# Divergent evolution of the Wnt signaling system in flatworms

**DOI:** 10.64898/2026.07.29.741434

**Authors:** Ludwik Gąsiorowski, Animan Tripathi, Elham Bavafaye Haghighi, Jochen C. Rink

**Affiliations:** Department of Tissue Dynamics and Regeneration, Max Planck Institute for Multidisciplinary Sciences, Göttingen, Germany; Institute of Evolutionary Biology, Faculty of Biology, University of Warsaw, Warsaw, Poland; International Max Planck Research School for Molecular Biology, Georg-August-University Göttingen, Göttingen, Germany; Faculty of Biology and Psychology, Georg-August-University Göttingen, Göttingen, Germany

## Abstract

Regenerative capacity varies widely across flatworms (Platyhelminthes). Whereas catenulids, microstomids and planarians can regenerate a complete head de novo, other flatworms cannot. This striking diversity raises a longstanding evolutionary question: does whole-body regeneration represent an ancestral trait that was subsequently lost in multiple lineages, or did it evolve convergently? Addressing this question requires comparative analyses of the molecular mechanisms underlying regeneration across phylogenetically diverse flatworms. Here, we focus on Wnt signaling, a deeply conserved regulator of antero-posterior (A–P) patterning and a central determinant of head-versus-tail identity during planarian regeneration, to establish a mechanistic framework for such comparisons. Although Wnt signaling has been studied extensively in planarians and parasitic neodermatans, its evolution and deployment in other flatworm clades remain poorly characterized. To address this gap, we characterized the complement of Wnt signaling components in two early-diverging flatworm clades, Catenulida and Macrostomorpha, with particular emphasis on expression and function in the catenulid *Stenostomum brevipharyngium*. Phylogenetic analyses reveal the ancient loss of six Wnt families and one secreted Frizzled-related protein (sFRP) family in the last common ancestor of flatworms, followed by additional lineage-specific gene losses and expansions. Moreover, several Wnt pathway components display markedly divergent expression patterns between catenulids and other flatworms, while functional analyses indicate corresponding differences in their regenerative deployment. Together, our findings reveal a dynamic evolutionary history of the flatworm Wnt signaling toolkit and establish a comparative framework for testing whether the molecular circuitry underlying head regeneration is ancestrally conserved or has evolved independently in distinct flatworm lineages.

## Introduction

Flatworms are famous for their whole-body regeneration abilities that allow replacement of both head and tail in bisected animals (Egger et al., 2007; Ivankovic et al., 2019; Newmark & Alvarado, 2002; Umesono et al., 2011). However, whole-body regeneration has been documented in only three flatworm clades (Tricladida, Macrostomorpha, and Catenulida), posing an important question: Is head regeneration an ancestral trait subsequently lost in many lineages, or did it evolve independently in each clade? Comparative studies of the molecular mechanisms of regeneration are crucial to answering this longstanding question (Srivastava, 2021).

The molecular underpinnings of head regeneration have been studied in detail in planarians, owing to the development of the two molecularly tractable model species, *Schmidtea mediterranea* and *Dugesia japonica* (Ivankovic et al., 2019; Newmark & Alvarado, 2002; Umesono et al., 2011). Upon wounding, planarians activate immediate early response genes (including paralogs of *runt*, *egr*, *jun* and *fos*) (Wenemoser et al., 2012), followed by the expression of Wnt1 at any wound site (Almuedo-Castillo et al., 2012; Gurley et al., 2010; Petersen & Reddien, 2008). Only anterior-facing wounds then activate the expression of the Wnt-inhibitor *notum* (Petersen & Reddien, 2008), which consequently mediates the wound-orientation selective inhibition of canonical Wnt signaling that is necessary and sufficient for head regeneration. Conversely, the activation of Wnt signaling (Gurley et al., 2008; Iglesias et al., 2008; Petersen & Reddien, 2008, 2011; Yazawa et al., 2009) and its subsequent maintenance via Wnt-dependent Wnt expression (Stückemann et al., 2017) is necessary for tail regeneration. Many Wnt signaling components are expressed in the body wall muscle in tail-to-head graded patterns in both regenerating and non-regenerating animals (Gurley et al., 2010). The resulting antero-posterior gradient of Wnt signaling activity maintains A-P axis identity during tissue turnover, not only in planarians (Gurley et al., 2010; Stückemann et al., 2017), but also in parasitic neodermatans (Armstrong et al., 2025; Jarero et al., 2024). Wnt signaling inhibitors (*sFRPs*, *notum*) are prominently expressed in the head and contribute to the maintenance and regeneration of the planarian A-P pattern (Gurley et al., 2010; Petersen & Reddien, 2008, 2011; Stückemann et al., 2017). Interestingly, the ability to regenerate the head is evolutionarily labile within planarians (Vila-Farré et al., 2023), and can be restored in several regeneration-deficient species via the inhibition of canonical Wnt signaling (Liu et al., 2013; Sikes & Newmark, 2013; Umesono et al., 2013; Vila-Farré et al., 2023).

Much less is known about the molecular control of regeneration in macrostomorphs and catenulids. In the former clade, whole-body regeneration has been documented so far only within a single family, Microstomidae (Palmberg, 1986), while in Catenulida, head regeneration abilities are widespread and reported in all major lineages (da Rosa & Loreto, 2022; Dirks et al., 2012; Gąsiorowski et al., 2025; Moraczewski, 1977; Van Cleave, 1929). So far, hardly any information on the molecular mechanisms of head regeneration in macrostomoprhs or catenulids exists, precluding direct comparison with planarians and answering the question of whether platyhelminth head regeneration is ancestral or convergently evolved.

To address this gap, we investigated the evolution and deployment of Wnt signaling in early-branching, non-model flatworms, with a particular focus on the catenulid *Stenostomum brevipharyngium*. First, we characterized the complement of Wnt ligands, receptors, and modulators in two catenulid and two macrostomorph species, generating the most comprehensive phylogenetic analysis of flatworm Wnt signaling components to date. We next determined the expression patterns of selected Wnt pathway components in intact *S. brevipharyngium* using hybridization chain reaction RNA in situ hybridization. Finally, we examined the expression dynamics and regenerative functions of selected Wnt signaling components during anterior and posterior regeneration through a combination of transcriptome sequencing, RNA in situ hybridization, and RNA interference. Together, our analyses reveal a dynamic evolutionary history of the flatworm Wnt signaling toolkit and substantial divergence in the expression and regenerative deployment of individual pathway components. Beyond these findings, they establish an essential comparative framework for testing whether the molecular circuitry underlying head regeneration is ancestrally conserved or has evolved independently in distinct flatworm lineages.

## Results

### Components of the Wnt signaling pathway among flatworms

So far, systematic analyses of flatworm Wnt signaling components have been performed only in planarians and neodermatans (Armstrong et al., 2025; Gurley et al., 2010; Jarero et al., 2024; Koziol et al., 2016; Riddiford & Olson, 2011). To gain a broader phylogenetic perspective on the evolution of Wnt ligands, their receptors (Frizzled proteins) and Frizzled-related antagonists (sFRPs), we mined the transcriptomes of two catenulids (*Stenostomum brevipharyngium* and *Stenostomum leucops*) and the genomes of two macrostomorphs (*Macrostomum cliftonense* and *Macrostomum hystrix*), which collectively represent two of the earliest-branching clades in flatworm phylogeny (Egger et al., 2015; Laumer et al., 2015). Putative Wnt signaling component orthologs were assigned to specific protein families based on maximum likelihood phylogeny inference of the encoded protein sequences (Figs. 1A and B, S1, S2).

**Fig. 1.**
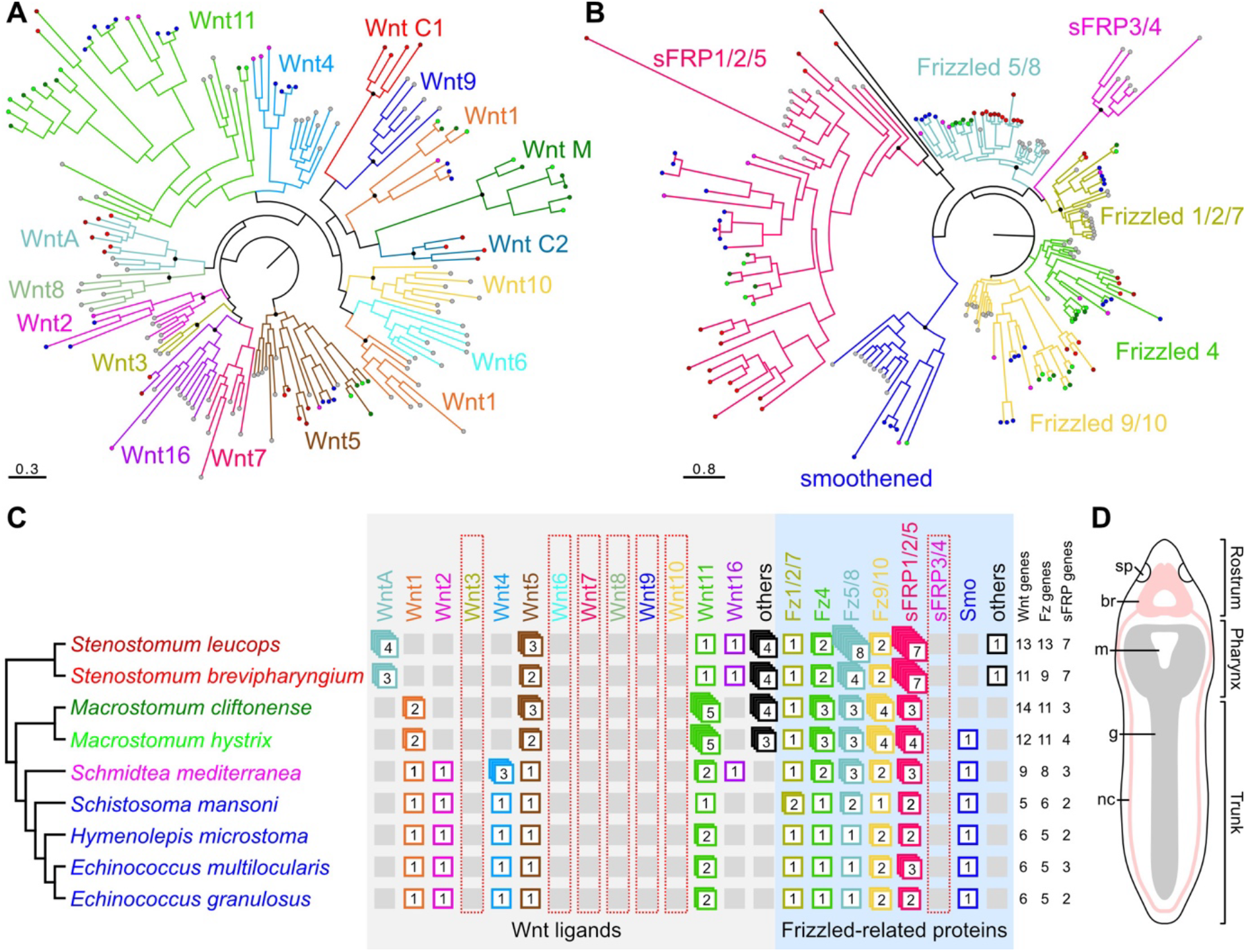
Evolution of Wnt signaling in flatworms. **A.** Maximum likelihood phylogeny of Wnt ligands. **B.** Maximum likelihood phylogeny of frizzled receptors and frizzled-related proteins. **C.** Complement of Wnt signaling components across platyhelminth phylogeny. **D.** Schematic drawing of *Stenostomum brevipharyngium,* used as a model catenulid in this study. Colored dots at the terminals in A and B represent sequences from the respective species of flatworms (same colors as in panel C); black dots at the nodes indicate a bootstrap value greater than 90%. Trees with full names of the terminals and all support values can be found as Figures S1 and S2. Abbreviations: *br* brain, *g* gut, *m* mouth opening, *nc* nerve cord, *sp* sensory pit.

For Wnt ligands, our analysis revealed striking differences in the conservation of particular bilaterian Wnt families between *Stenostomum* and *Macrostomum*, with only minor differences in paralog numbers within each genus (Fig. 1A, C; Tab. 1). Wnt3 appears to be absent in all the analyzed flatworms, and WntA is only present in *Stenostomum*. Both *Stenostomum* species have multiple copies of WntA and Wnt5 and single copies of Wnt11 and Wnt16, but their transcriptomes do not include the other canonical Wnt ligands. Moreover, in each species of *Stenostomum,* we found four Wnt ligands that could not be grouped into known Wnt families and instead form two independent catenulid-specific groups (we called them WntC1 and WntC2). Both species of *Macrostomum* have two copies of Wnt1, multiple copies of Wnt5 and Wnt11, and multiple copies of the macrostomorph-specific Wnt ligands (WntM), with the other Wnt ligands missing. Our systematic phylogeny informs the classification of *S. mediterranea* Wnt orthologs (Fig. 1C, Tab. 1), which remain ambiguous in some cases. Consistent with the previous results of Riddiford and Olson (2011), out of the six Wnt paralogs initially designated as Wnt11 by Gurley et al. (Gurley et al., 2010), three actually cluster with Wnt4 (Wnt11-4, −5 and 6); one (Wnt11-3/Wnt-P-4) confidently clusters with the Wnt16 group, while only two remain in the Wnt11 group (Wnt11-1 and Wnt11-2) (Fig. 1A; Table 1). Consequently, our analysis indicates planarians have single orthologs of Wnt1, Wnt2, Wnt5 and Wnt16, three paralogs of Wnt4, and two paralogs of Wnt11. In Neodermata, there are single orthologs of Wnt1, Wnt2, Wnt4, and Wnt5 and two paralogs of Wnt11 (with the exception of *Schistosoma mansoni*, which has only a single Wnt11 paralog). Similarly to the previous results (Riddiford & Olson, 2011), platyhelminth Wnt1 proteins do not group with the orthologous sequences from other Bilateria (Fig. 1A). Altogether, six bilaterian Wnt families (Wnt3, Wnt6, Wnt7, Wnt8, Wnt9, and Wnt10) seem to be missing in all flatworms examined, indicating loss of these genes in the common ancestor of platyhelminths (Fig. 1C), while the subsequent losses and duplications occurred independently in particular flatworm ingroups.

**Table 1.** Orthology and nomenclature of Wnt genes in Rhabditophora and Catenulida (here represented by a planarian *Schmidtea mediterranea* and a catenulid *Stenostomum brevipharyngium,* respectively).

| subfamily | Rhabditophora |  |  |  | Catenulida |  |
| --- | --- | --- | --- | --- | --- | --- |
|  |  | Riddiford<br>and Olson<br>(2011) | Gurley et<br>al. (2010) | alternative<br>names | this study |  |
| WntA | absent |  |  |  | present | WntAa<br>WntAb<br>WntAc |
| Wnt1 | present | Wnt1 | Wnt1 | WntP1 | absent | - |
| Wnt2 | present | Wnt2 | Wnt2 | - | absent | - |
| Wnt3 | absent | - | - | - | absent | - |
| Wnt4 | present | Wnt4a<br>Wnt4b<br>Wnt4c | Wnt11-4<br>Wnt11-5<br>Wnt11-6 | WntP3<br>WntP2<br>WntA | absent | - |
| Wnt5 | present | Wnt5 | Wnt5 | - | present | Wnt5a<br>Wnt5b |
| Wnt6 | absent | - | - | - | absent | - |
| Wnt7 | absent | - | - | - | absent | - |
| Wnt8 | absent | - | - | - | absent | - |
| Wnt9 | absent | - | - | - | absent | - |
| Wnt10 | absent | - | - | - | absent | - |
| Wnt11 | present | Wnt11a<br>Wnt11b | Wnt11-1<br>Wnt11-2 | - | present | Wnt11 |
| Wnt16 | present | WntP4 | Wnt11-3 | WntP4 | present | Wnt16 |
| Non-canonical<br>Wnts |  |  |  |  | present | WntC1a<br>WntC1b<br>WntC2a<br>WntC2b |

Frizzled receptors from all major families (Fz1/2/7, Fz4, Fz5/8 and Fz 9/10) are present in at least a single copy in each analyzed flatworm species (Fig. 1B, C). Most of them, except Fz1/2/7, occur in multiple copies in free-living flatworms (with the record number of eight copies of Fz5/8 in *S. leucops*), while only sporadic duplications are present in neodermatans. One of the two canonical families of sFRPs (sFRP3/4) is entirely missing in all flatworms, indicating an ancient loss of the gene in the platyhelminth lineage (Fig. 1C). In contrast, the other family, sFRP1/2/5, has likely undergone expansion, especially in *Stenostomum*, in which it includes seven copies. Additionally, both species of *Stenostomum* have a single sFRP-like gene that could not be confidently placed in any of the families of Frizzled-like proteins, but instead groups with the sFRP1/2/5 family. Finally, *Smoothened*, a phylogenetically related protein that is involved in hedgehog and not Wnt signaling, is present as a single ortholog in most flatworm species, but it is missing in both species of *Stenostomum* and in *M. cliftonense* (Fig. 1C).

### Expression of Wnt signaling components in catenulids

Next, we set out to study the expression patterns of Wnt signaling components in a catenulid, *S. brevipharyngium* (Fig. 1D). The three WntA paralogs are generally expressed in the anterior structures (Fig. 2A). WntAa is expressed in distinct neurons in the brain (Fig. 2B) and in a few cells in front of the mouth opening (Fig. 2C). WntAb and WntAc have strongly overlapping domains that include some neurons in the posterior brain (Fig. 2B) and cells in the anterior part of the pharynx (Fig. 2C).

**Fig. 2.**
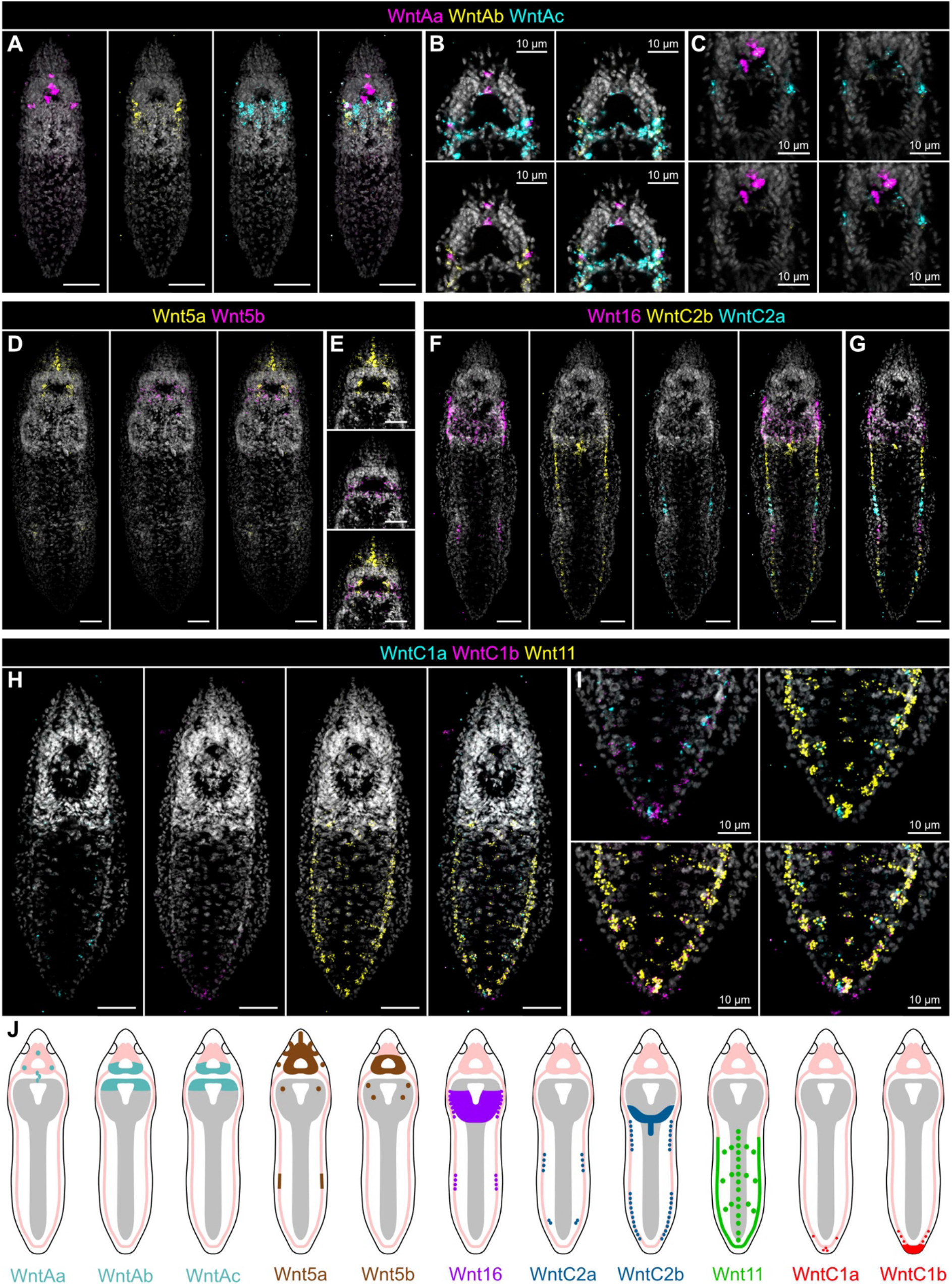
Expression of Wnt ligands in a catenulid *Stenostomum brevipharyngium*. **A – I.** in situ RNA hybridization chain reactions. Names of the hybridized genes are provided above the panels. Cell nuclei are counterstained with Hoechst (grey). Scale bars represent 20 μm if not stated otherwise. **J.** Schematic drawings summarizing the expression of particular Wnt ligands.

The two Wnt5 paralogs are also mostly restricted to the anterior region (Fig. 2D). Wnt5a is expressed in the brain (where it is co-expressed with Wnt5b), in a stripe of cell in the middle of the rostrum, and in the sensory pits (Fig. 2E). Additionally, faint Wnt5a expression can be observed in the middle of the trunk (Fig. 2D). Wnt5b displays a more restricted expression domain in the mid-posterior brain neurons (Fig. 2D, E).

Wnt16, WntC2a, and WntC2b show interesting complementary patterns along the A-P axis (Fig. 2F, G). Wnt16 is expressed in the pharynx, behind the mouth opening, and in paired lateral domains along the pharynx, and also more posteriorly in the mid-trunk. WntC2b is expressed in the most posterior region of the pharynx and a few medio-ventral cells just behind the pharynx. Additionally, WntC2b is expressed in lateral domains that overlap with the posterior parts of the Wnt16 lateral domains but extend more posteriorly, both in the pharyngeal region and in the mid-trunk. Finally, WntC2a is also expressed in the lateral domains, overlapping with the posterior expression of WntC2b and extending even more posteriorly. Consequently, the lateral domains include regions that are Wnt16^+^/WntC2b^-^ /WntC2a^-^, Wnt16^+^/WntC2b^+^/WntC2a^-^, Wnt16^-^/WntC2b^+^/WntC2a^-^, Wnt16^-^ /WntC2b^+^/WntC2a^+^, and finally Wnt16^-^/WntC2b^-^/WntC2a^+^. This pattern is repeated once, starting from the middle of the trunk (Fig. 2G).

The single ortholog of Wnt11 is broadly expressed in lateral domains, along the entire trunk of the animal (Fig. 2H). Additionally, the gene is expressed on the ventral side of the worm, both in the midventral stripe and in cells that form a repetitive pattern of inverted-V-shaped domains (Fig. 2H). This results in a high concentration of Wnt11^+^ cells at the posterior pole (Fig. 2I).

Finally, two WntC1 paralogs are restricted to the most posterior structures (Fig. 2H). WntC1a is expressed in some of the most posterior Wnt11^+^ cells (Fig. 2I). WntCb is expressed in another, only partially overlapping set of the posterior Wnt11^+^ cells but also in epidermal cells at the very posterior tip of the worm that do not express Wnt11 (Fig. 2I). Therefore, there are sets of posterior cells with five different combinations of Wnt expression: Wnt11^+^/WntC1a^-^ /WntC1b^-^, Wnt11^+^/WntC1a^+^/WntC1b^-^, Wnt11^+^/WntC1a^-^/WntC1b^+^, Wnt11^+^/WntC1a^+^/WntC1b^+^, and Wnt11^-^/WntC1a^-^/WntC1b^+^.

In summary, Wnt ligands are expressed throughout the entire body of *S. brevipharyngium*, from the rostrum to the tip of the tail (Fig. 2J). Based on their specific expression patterns, the Wnt ligands can be split into an anterior-restricted group (WntA, Wnt5), a trunk-specific group expressed in repeating stripe-like patterns (Wnt16, WntC2), and a posterior group that is restricted to the posterior trunk (Wnt11, WntC1). Interestingly, we failed to observe both tail-to-head graded expression patterns as prominently displayed by Smed-Wnt11-1 or Smed-Wnt11-5/P-2, or prominent tail tip expression as displayed by the tail pole marker Smed-Wnt1, suggesting that Wnt expression in *Stenostomum* is qualitatively different from *S. mediterranea* (see discussion).

We similarly analyzed the expression of Frizzled receptors and sFRPs in intact worms (Fig 3). Fz1/2/7 is broadly expressed from the posterior brain to the mid-trunk (Fig. 3A). Particularly strong and uniform expression can be detected in the pharynx. Fz4a is expressed anteriorly in the rostrum and just behind the pharynx (Fig. 3B), while expression of Fz4b is restricted to the posterior part of the pharynx (Fig. 3C). Four paralogs of Fz5/8 are generally expressed in the anterior part of the animal (3D–G). Fz5/8a and Fz5/8b are expressed in the anterior tip of the worm, in front of the sensory pits (Fig. 3D, E). Fz5/8c is expressed in the central brain and in the rostrum (Fig. 3F), while Fz5/8d^+^ cells are scattered throughout the head and pharyngeal regions (Fig. 3G). Two paralogs of Fz9/10 show similar, weak expression across most of the trunk (Fig. 3H–I).

**Fig. 3.**
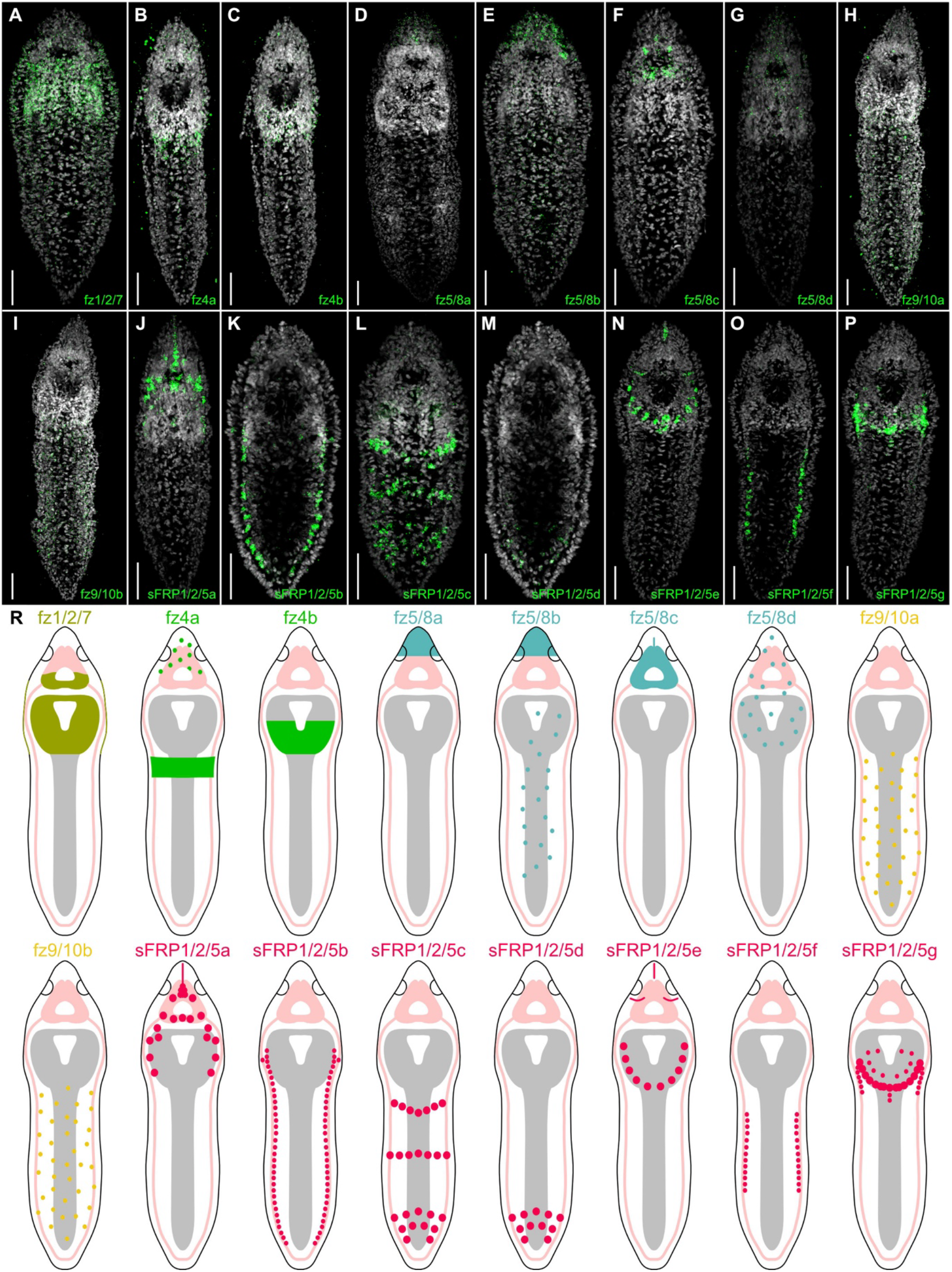
Expression of frizzled receptors and secreted frizzled-related proteins (sFRPs) in the catenulid *Stenostomum brevipharyngium*. **A – P.** in situ RNA hybridization chain reactions. Names of the hybridized genes are provided in the lower-right corner of each panel. Cell nuclei are counterstained with Hoechst (grey). Scale bars represent 20 μm. **R.** Schematic drawings summarizing the expression of particular genes.

Seven paralogs of sFRP1/2/5 show very distinct and divergent expression patterns (Fig. 3J–P). sFRP1/2/5a is expressed in a stripe of cells in the middle of the rostrum, in some brain neurons, and in scattered cells in the pharynx (Fig. 3J). sFRP1/2/5b is expressed throughout the entire trunk in lateral domains (Fig. 3K). sFRP1/2/5c has three transverse expression domains in the anterior, mid, and posterior trunk (Fig. 3L). sFRP1/2/5d is only restricted to the posterior domain in the trunk (Fig. 3M). sFRP1/2/5e is expressed in the middle of the rostrum and in the ring of cells surrounding the mouth opening (Fig. 3N). sFRP1/2/5f is expressed in the same lateral domains as sFRP1/2/5b, but only in the middle portion of the trunk, while it remains absent from the anterior and posterior extremities of the trunk (Fig. 3O). Finally, sFRP1/2/5g is predominantly expressed in the posterior pharynx, but also in some anterior cells in the lateral domains (Fig. 3P).

In general, Frizzled paralogs show similar, staggered expression along the antero-posterior axis, with paralogs of Fz5/8 predominantly expressed in the anterior, Fz1/2/7 and paralogs of Fz4 in the pharynx and anterior trunk, and paralogs of Fz9/10 in the trunk (Fig. 3R). In contrast, seven paralogs of SFRP1/2/5 show divergent expression patterns spanning the entire body axis of *S. brevipharyngium* (Fig. 3R).

### Expression of Wnt signaling components in muscles

In planarians and neodermatans, Wnt pathway components are predominantly expressed within body muscle (Jarero et al., 2024; Witchley et al., 2013). We aimed to test if this is also the case in Catenulida. The musculature of catenulids can be detected by phalloidin staining and is composed of five major components: rostral musculature (including longitudinal, circular and helical muscles), pharyngeal musculature (including longitudinal and circular muscles of the pharyngeal bulb and pharyngeal dilators that connect the pharyngeal bulb with the body walls), longitudinal muscles of the trunk, circular muscles of the trunk and dorso-ventral muscles of the trunk (Fig. 4A and B).

**Fig. 4.**
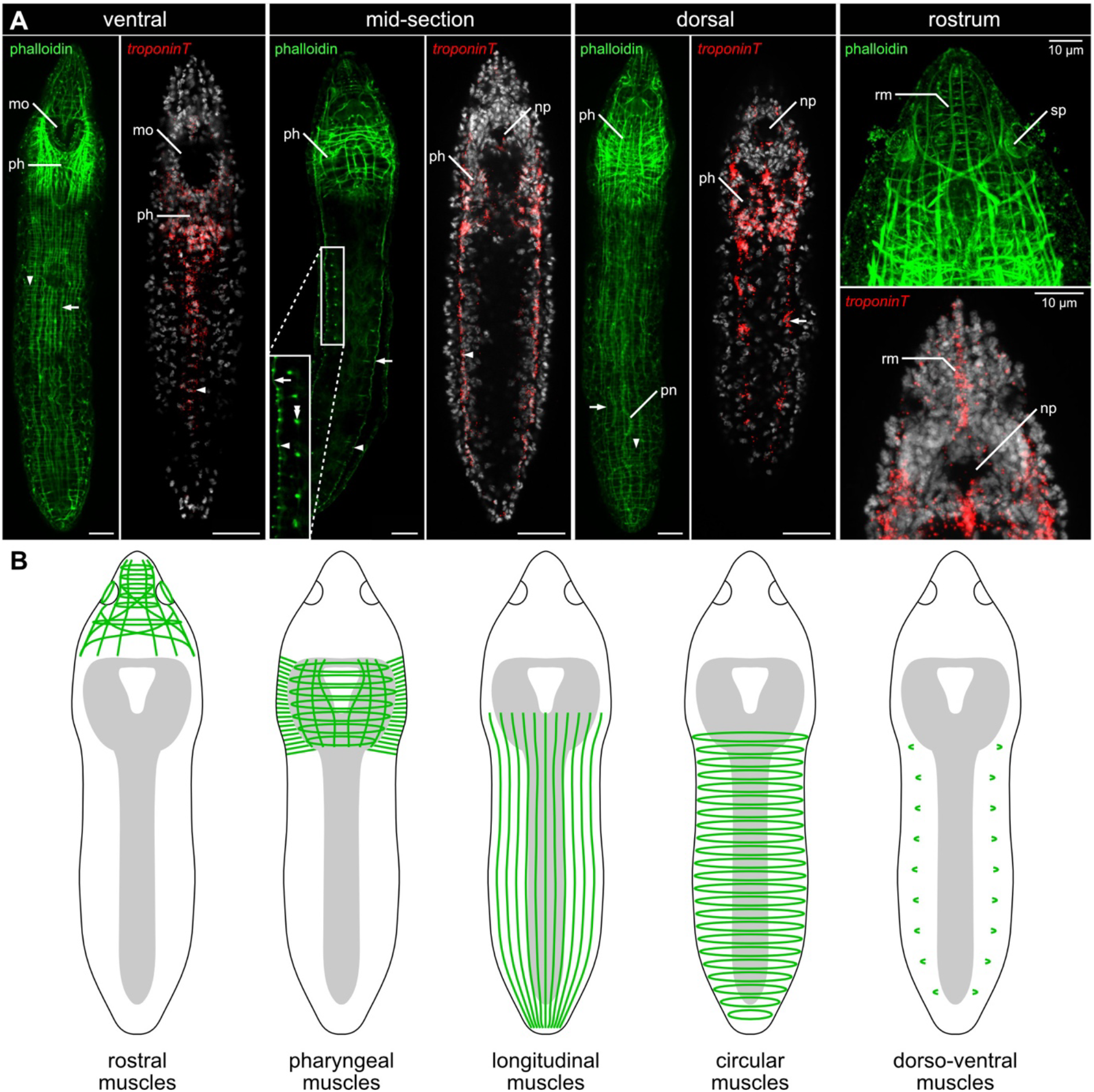
Musculature of *Stenostomum brevipharyngium*. **A.** Musculature in different sections of the body visualized with phalloidin (green) and in situ RNA hybridization chain reaction against muscle marker *troponinT* (red). Cell nuclei are counterstained with Hoechst (grey). Scale bars represent 20 μm if not stated otherwise. Arrowheads indicate circular muscles, arrows longitudinal muscles, and double arrowheads dorso-ventral muscles. Abbreviations: *mo* mouth opening, *np* brain neuropile, *ph* pharynx, *pn* protonephridium, *rm* rostral muscles, *sp* sensory pit. **B.** Schematic drawings of different portions of musculature.

Unfortunately, phalloidin staining is incompatible with the HCR in situ protocol that we used to visualize gene expression. To overcome this limitation, we used expression of the muscle cell marker *troponinT* (Gąsiorowski et al., 2025) to visualize nuclei of the muscle cells, which in some cases can be directly associated with particular sets of muscles (Fig. 4A). The nuclei of rostral muscles form a distinct line in the middle of the rostrum. Most of the cells in the pharyngeal and buccal region are *troponinT*^+^, representing muscular components of the pharynx and mouth apparatus. Additionally, there are densely packed *troponinT* ^+^ cells located laterally and ventrally in the trunk (which likely represent nuclei of the circular muscles). Additional *troponinT* ^+^ cells scattered throughout the trunk correspond to the nuclei of the longitudinal muscles.

First, we concentrated on the expression of several genes that are expressed in a stripe of cells in the middle of the rostrum (Fig. 5). Those cells express Wnt5a, sFRP1/2/5a, and Fz5/8c and are positive for *troponinT*, indicating they are rostral myocytes (Fig. 5A – C). Interestingly, the same set of cells also expresses two *Notum* paralogs that show further weak expression in the pharyngeal region (Fig. 5D and E). Next, we focused on the cells in the lateral domains that show specific combinations of the expression of particular Wnt signaling components along A-P axis (Fig. 6). The lateral cells expressing WntC2a, WntC2b, sFRP1/2/5b and sFRP1/2/5f were all *troponinT*^+^, suggesting that these genes might be involved in patterning the A-P identity of the trunk circular muscles (Fig. 6A – C). We also found that WntC2b^+^ cells in the mid-ventral trunk (Fig. 6E) and sFRP1/2/5e^+^ and sFRP1/2/5f^+^ cells in the pharynx and in the trunk all co-express *troponinT* (Fig. 6F – J). In summary, all the tested Wnt signaling components were co-expressed with *troponinT*, indicating these genes are predominantly expressed by the muscular system.

**Fig. 5.**
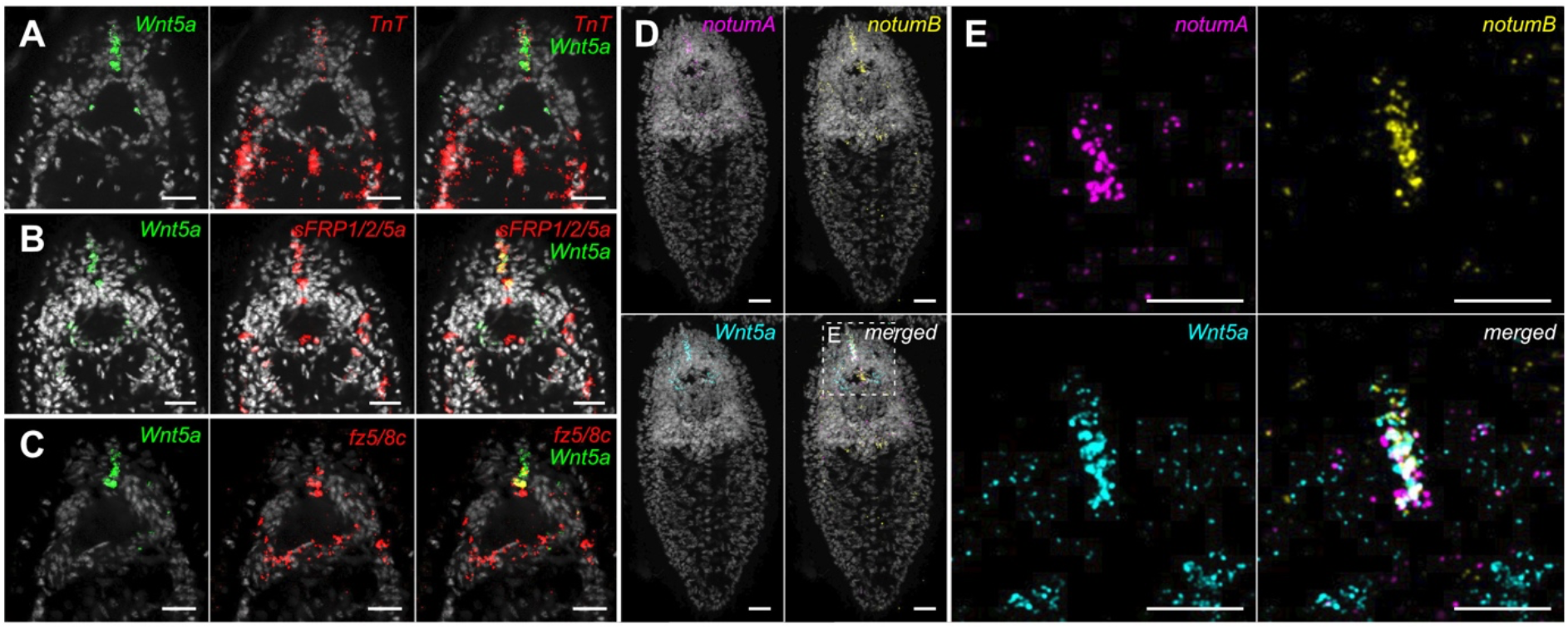
Gene expression in the rostral muscular domain of *Stenostomum brevipharyngium*. **A – C**, co-expression of *Wnt5a*, *troponin T*, *sFRP1/2/5a* and *fz 5/8c*. **D – E** co-expression of *Wnt5a* notum A and B. Cell nuclei are counterstained with Hoechst (grey). Scale bars represent 10 μm. The box in panel D indicates the area magnified in panel E.

**Fig. 6.**
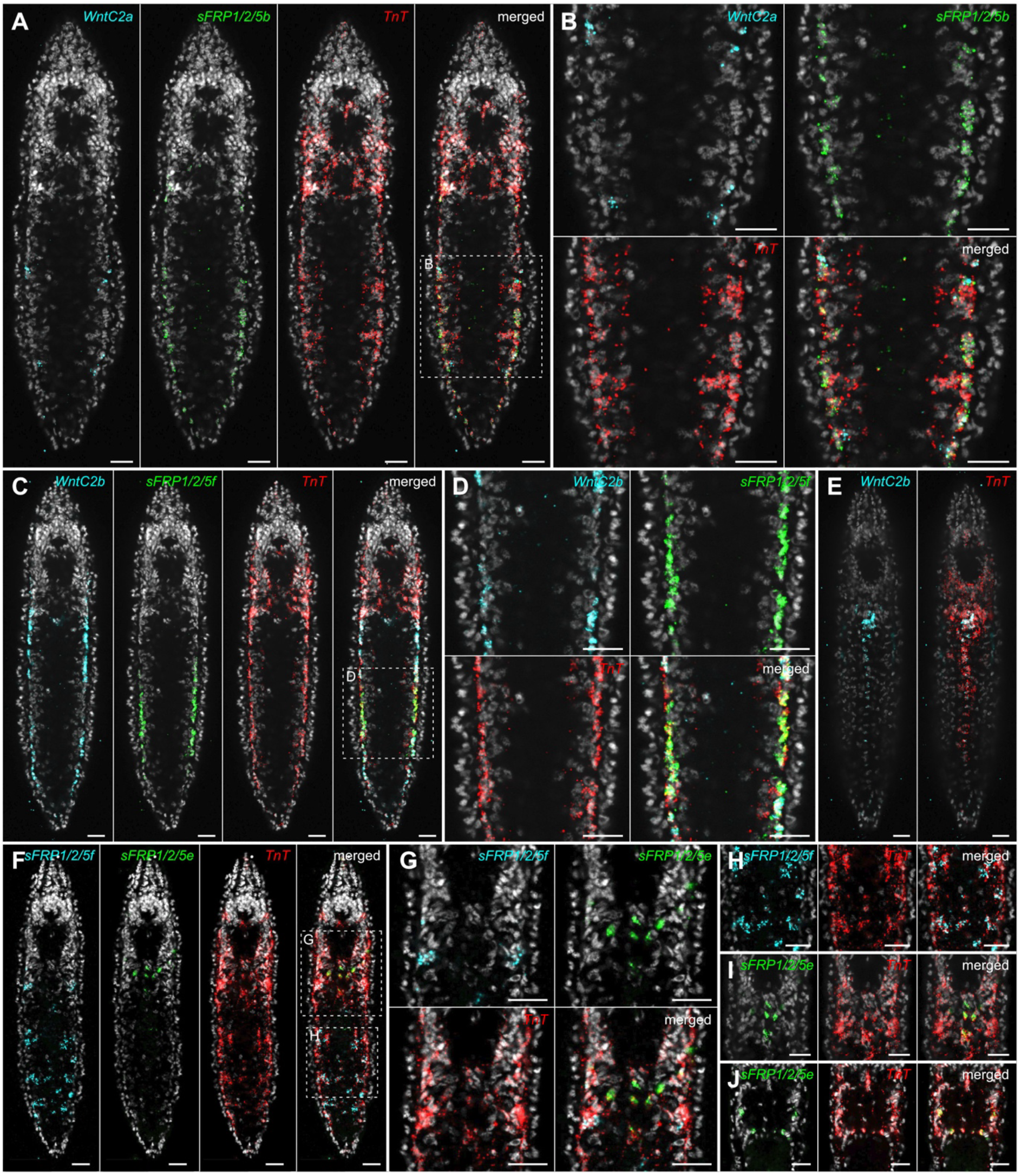
Components of the Wnt signaling pathway are expressed by muscle cells in *Stenostomum brevipharyngium,* as indicated by co-expression with the muscle marker *troponin T* (red). Names of the hybridized genes are provided in the upper-right corner of each panel. Cell nuclei are counterstained with Hoechst (grey). Scale bars represent 10 μm. Boxes on panels A, C, and F indicate the areas magnified in the respective labelled panels.

*Gene expression dynamics during regeneration in* S. brevipharyngium

To study the temporal dynamics of Wnt pathway component expression during regeneration, we performed time-course RNA sequencing of worms at different stages of anterior and posterior regeneration (Fig. 7A). The worms were either cut through the pharynx to initiate head regeneration or the posterior tip was amputated to elicit tail regeneration. 60 worms were cut and pooled for each sample, and total RNA was extracted at 0, 1, 5, 12, 24, and 48 hours post amputation (hpa). Three independently generated RNA replicates were sequenced per regeneration time point and regeneration paradigm, and an additional three sets of 60 intact worms were sequenced as a baseline reference.

**Fig. 7.**
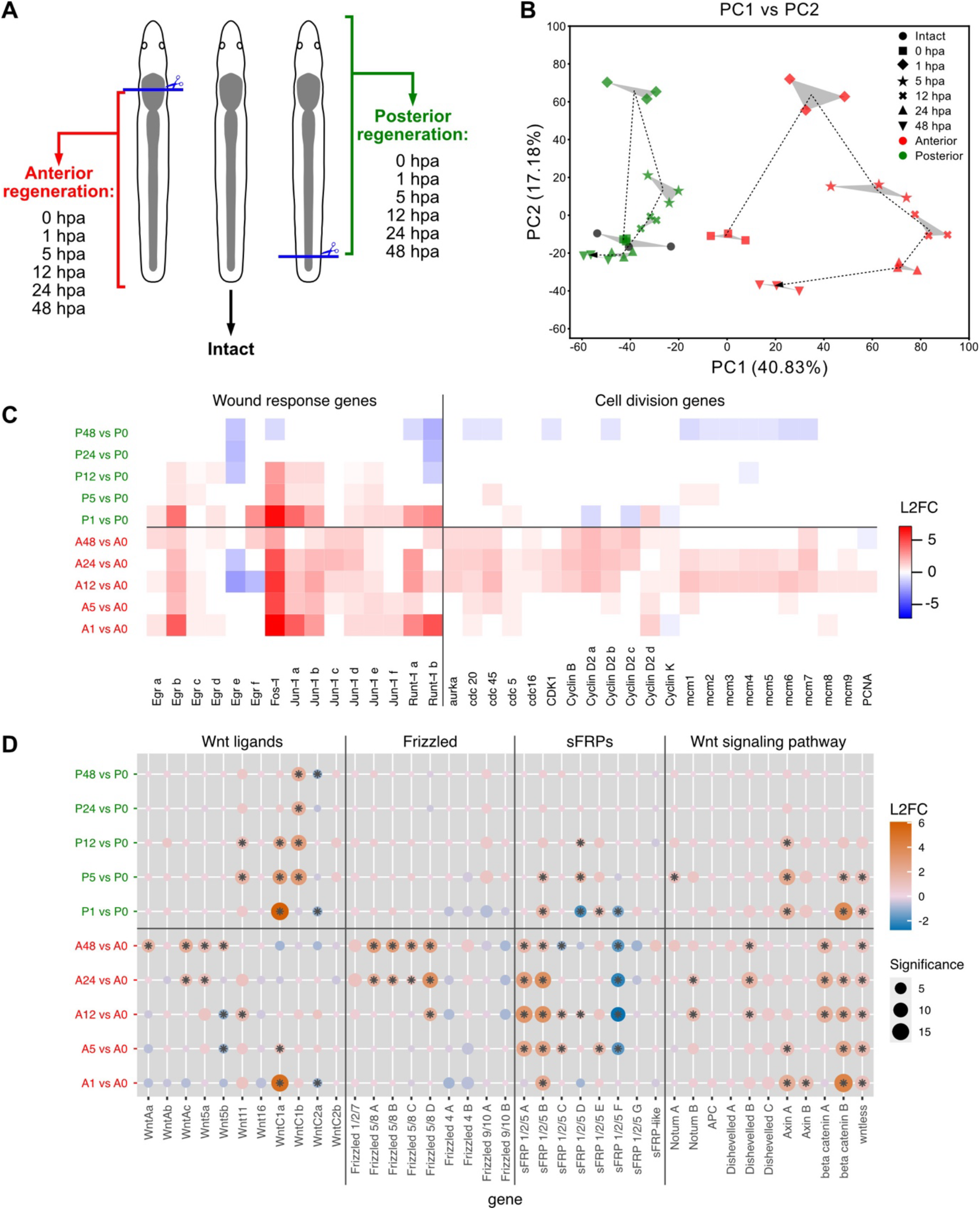
Transcriptomic response during anterior and posterior regeneration of *Stenostomum brevipharyngium*. **A.** Amputation paradigm and experimental design. **B.** Principal component analysis of sequenced samples. Dotted lines show the progression of anterior (red) and posterior (green) regeneration temporal trajectories. **C.** Heatmap of significant (adjusted p-value <0.05) differences in the expression of wound response and cell division genes during anterior and posterior regeneration. **D.** Dot plot showing significance and log2fold change value for the Wnt signaling components during anterior and posterior regeneration. Abbreviations: A0–A48 anterior (head) regeneration at 0–48 hpa, P0–P48 posterior (tail) regeneration at 0–48 hpa.

We first performed principal component analysis of the transcriptomic data (Fig. 7B). ∼58% of the total variation in gene expression was explained by the first two principal components, and the close grouping of the replicates at each time point indicated a strong signal-to-noise ratio. Consistently, both the anterior and posterior time series followed distinct trajectories through PCA-space. For anterior regeneration, the 0 hpa samples grouped relatively close to the intact worms, but the strong displacement at 1 hpa indicates the existence of a strong wound-induced gene expression response. At 48 hpa, the samples were still clearly distinct from those from intact worms, in accordance with our previous morphology-based assessment showing that the completion of head regeneration in *S. brevipharyngium* takes about 96h (Gąsiorowski et al., 2025; Gąsiorowski, Dittmann, et al., 2023; Tratkiewicz & Gąsiorowski, 2025). For posterior regeneration, the wound-induced shift in gene expression between 0 hpa and 1 hpa was also clearly apparent, yet somewhat less pronounced than in the case of anterior regeneration. However, the tail regeneration gene expression trajectory converged back to that of intact worms at 24 hpa, indicating that tail regeneration is already largely complete at that time point, and thus much faster than head regeneration.

Next, we assessed the expression dynamics of canonical wound response genes (*Egr*, *Fos-1*, *Jun-1*, and *Runt-1*) (Srivastava, 2021; Wenemoser et al., 2012) and various genes involved in cell division in *Stenostomum* (Gąsiorowski et al., 2025) during both anterior and posterior regeneration (Fig. 7C). Most of the wound response genes showed the expected patterns of strong up-regulation in the early stages of both head and tail regeneration, with a slow decay over time. Interestingly, the wound response seemed to be more pronounced and longer lasting during head regeneration (most of the genes still upregulated at 48 hpa), when compared to tail regeneration (upregulation only until 12 hpa). While these results agree with the observation that the process of anterior regeneration takes much longer than posterior regeneration (Fig. 7B), they might also reflect the proportionally greater loss of total cell mass during the head amputation paradigm. The head-regeneration-specific upregulation of cell division genes between 12-48 hpa (Fig. 7C) is consistent with a greater requirement for cell divisions to reconstitute the morphologically complex anterior structures (i.e., brain and pharynx), when compared to the tail that lacks distinct morphological structures.

Finally, we examined the expression dynamics of individual Wnt pathway components during tail and head regeneration. For both time series, we extracted differentially expressed (DE) genes versus the 0 hpa baseline (Fig. 7C). In general, Wnt components expressed in the tail of intact worms responded during posterior regeneration, while head-expressed components responded during anterior regeneration. For example, the tail-specific Wnts (WntC1b and Wnt11) became significantly upregulated by 5 h after tail removal. For the paralogs of the head-associated WntA and Wnt5 ligands, significant changes in expression only became apparent at the later stages of head regeneration (24–48 hpa), in accordance with their expression in the brain and rostral structures, which reappear in late stages of regeneration. However, the most dramatic regeneration-associated expression change of any Wnt ligand was observed for WntC1a, which is weakly expressed at the tail tip of intact animals, but becomes dramatically and transiently upregulated at the very early stages (1 hpa) of both head and tail regeneration. This finding is intriguing, as it parallels the similarly dramatic, early upregulation of planarian posterior Wnt1 at any wound site (Gurley et al., 2010; Petersen & Reddien, 2009).

In contrast to the Wnt ligands, the Frizzled receptors did not show significant expression changes during posterior regeneration, consistent with the generally very low expression of Fz receptors in the tail. During head regeneration, all head-specific paralogs of Frizzled 5/8 became upregulated between 12 and 24 hpa. The sFRPs generally responded more dynamically than the Frizzled receptors, showing similar patterns in both anterior and posterior regeneration, although with less pronounced shifts in the latter. The only exception was sFRP1/2/5A, which showed strong up-regulation during anterior regeneration, starting from 5hpa onwards, in accordance with its expression in the anterior structures. We also investigated transcriptional changes of other important canonical Wnt signal transduction components, including the paralogs of *notum*, *APC*, *Dishevelled*, *axins*, *β-catenin,* and *wintless*. Interestingly, *wintless* and at least one of the two *axin* paralogs were significantly upregulated already at the 1 h time point in both head and tail regeneration, reminiscent of their early upregulation after any wound in *S. mediterranea* (Stückemann et al., 2017). *notum B* was significantly upregulated at 12 and 24 hpa of head regeneration only, while *notum A* was significantly upregulated only at 5 hpa during tail regeneration. Both the low fold-change of

*notum B* expression and the tail regeneration specificity of *notum A* contrast with the pronounced, head-specific induction of the single *notum* homolog in *S. mediterranea*. *S. brevipharyngium* has two paralogs of *β-catenin* resulting from *Stenostomum*-specific duplication, independent of the expansion of *β-catenin* genes in other flatworms (Fig. S3A; Kibet et al., 2025). Both of those paralogs possess multiple Armadillo repeat domains (Fig. S3B), which are believed to be involved in Wnt signaling, and thus potentially might play a role in the regulation of the Wnt system. One of those paralogs, *β-catenin A*, is upregulated from 12 hpa onwards in the anteriorly regenerating worms, while *β-catenin B* shows strong upregulation at both anterior and posterior wounds already from 1hpa. The significant upregulation of the *S. brevipharyngium β-catenin B* already at 1 hpa of both head and tail regeneration is a further notable difference to *Schmidtea mediterranea*, where the canonical Wnt-signaling-associated Smed-β-catenin-1 is constitutively expressed and thought to be entirely regulated on a posttranscriptional level.

To complement the temporal dynamics of the RNA-seq gene expression data with spatial information, we next turned to whole-mount HCR experiments. Both WntC1a, the only wound-responsive Wnt paralog in *Stenostomum*, and the canonical wound-response gene *Runt-1A* displayed strong upregulation in both anteriorly and posteriorly regenerating worms at 1hpa (Fig. 8). In worms regenerating their head (Fig. 8A), WntC1a was strongly induced in a small number of cells in the pharyngeal region distal to the wound edge, and, to a lesser extent, in cells at discrete locations within the trunk. In contrast, *Runt-1A* was strongly and broadly expressed in cells immediately adjacent to the wound edge, with gradually decreasing expression extending into the trunk and tail regions. In tail-amputated animals (Fig. 8B), WntC1a was induced proximal to the amputation wound in several tiers of cells in the entire trunk, while *Runt-1A* was broadly expressed in the immediate wound vicinity and, with gradually decreasing expression, extending as far proximal as the rostrum. To probe the identity of the cells in which the wound-induced upregulation of WntC1a occurs, we next performed double in situ hybridization of WntC1a and the muscle marker *troponinT*. In both anteriorly and posteriorly amputated worms at 1 hpa, we detected strong and specific co-localization of gene expression (Fig. 8C-H). Altogether, these experiments indicate the wound-induced upregulation of WntC1a occurs in a subset of muscle cells, similar to the wound-induced expression of Wnt1 in the body wall musculature of *S. mediterranea*. However, WntC1a upregulation in *S. brevipharyngium* is not restricted to the immediate wound vicinity like *S. mediterranea* Wnt1, instead extending to cohorts of muscle cells distant to the wound site.

**Fig. 8.**
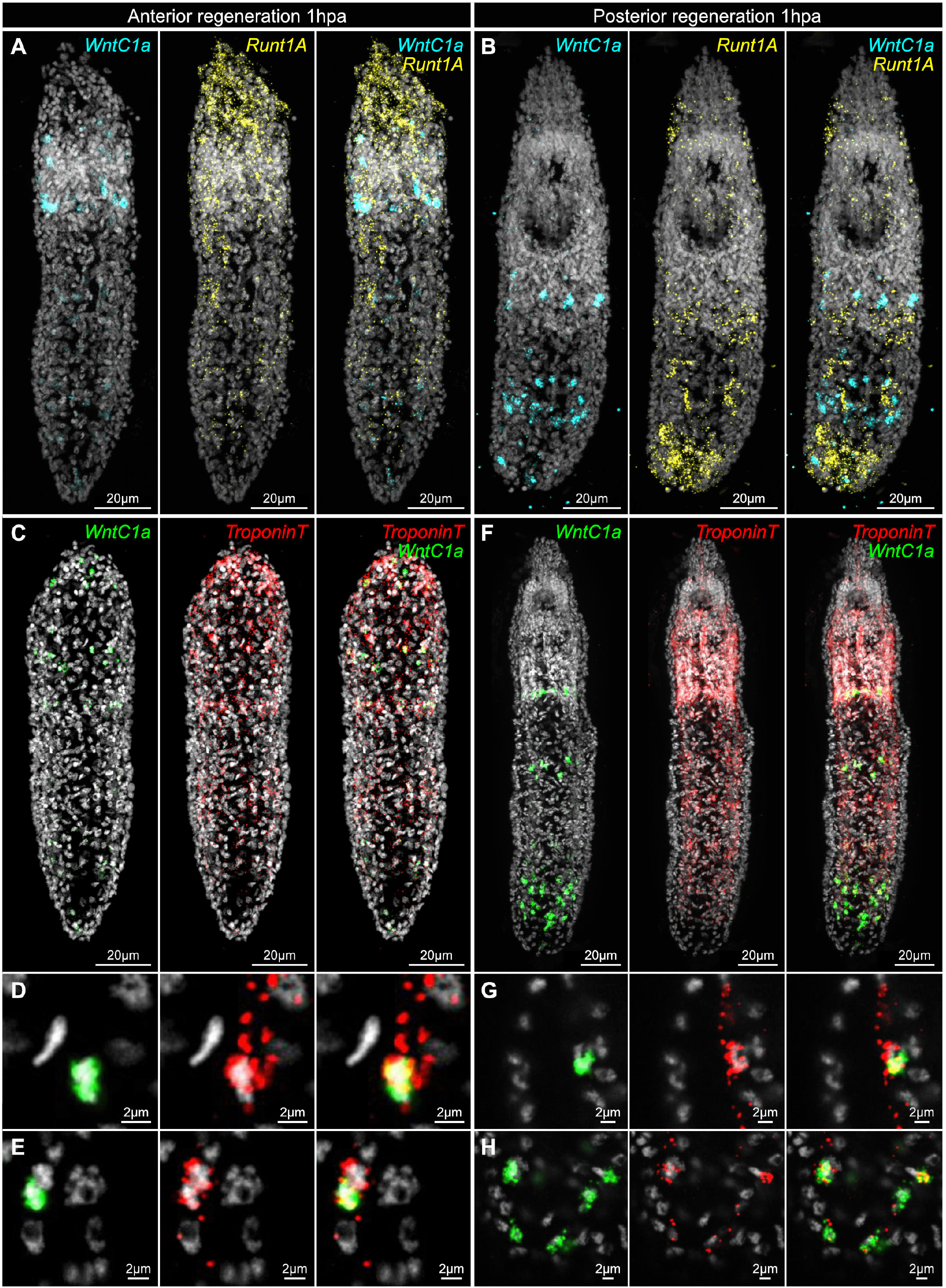
Wound-induced gene expression in *Stenostomum brevipharyngium* during head (**A, C, D, E**) and tail (**B, F, G, H**) regeneration at 1 hour post-amputation (hpa). Names of the hybridized genes are provided in the upper-right corner of each panel. Cell nuclei are counterstained with Hoechst (grey).

Next, we examined the expression patterns of the two *notum* paralogs during the early stages of head and tail regeneration in *S. brevipharyngium* (Fig. 9). Neither displayed striking changes at 1 and 5 h post head or tail amputation, consistent with the weak wounding response of the genes in the RNAseq data (Fig. 7D). At 1 hpa, the *notum* paralogs generally showed low expression in both head and tail regeneration (Fig. 9A and B), yet in the trunks of both regenerates, we could observe some *notum B*^+^ cells that are absent in the intact worms (compare Fig.9A and B with Fig. 5D). In contrast, at 5 hpa, both genes show stronger expression in the tissues distal from the wound site in both regeneration paradigms (Fig. 9C and D). In the head regenerating worms, both *notums* are expressed in the pharyngeal region and in the non-overlapping domains in the lateral trunk (Fig. 9C). In the tail regenerating animals, they are strongly expressed in the intact anterior domains, while *notum A* shows additional strong expression in the trunk (Fig. 9D), in accordance with our transcriptomic data that show up-regulation of *notum A* in the tail-regenerating worms at 5hpa (Fig. 7D). In general, these data are consistent with the absence of wound-induced expression of the two *notum* paralogs in our RNA-seq data.

**Fig. 9.**
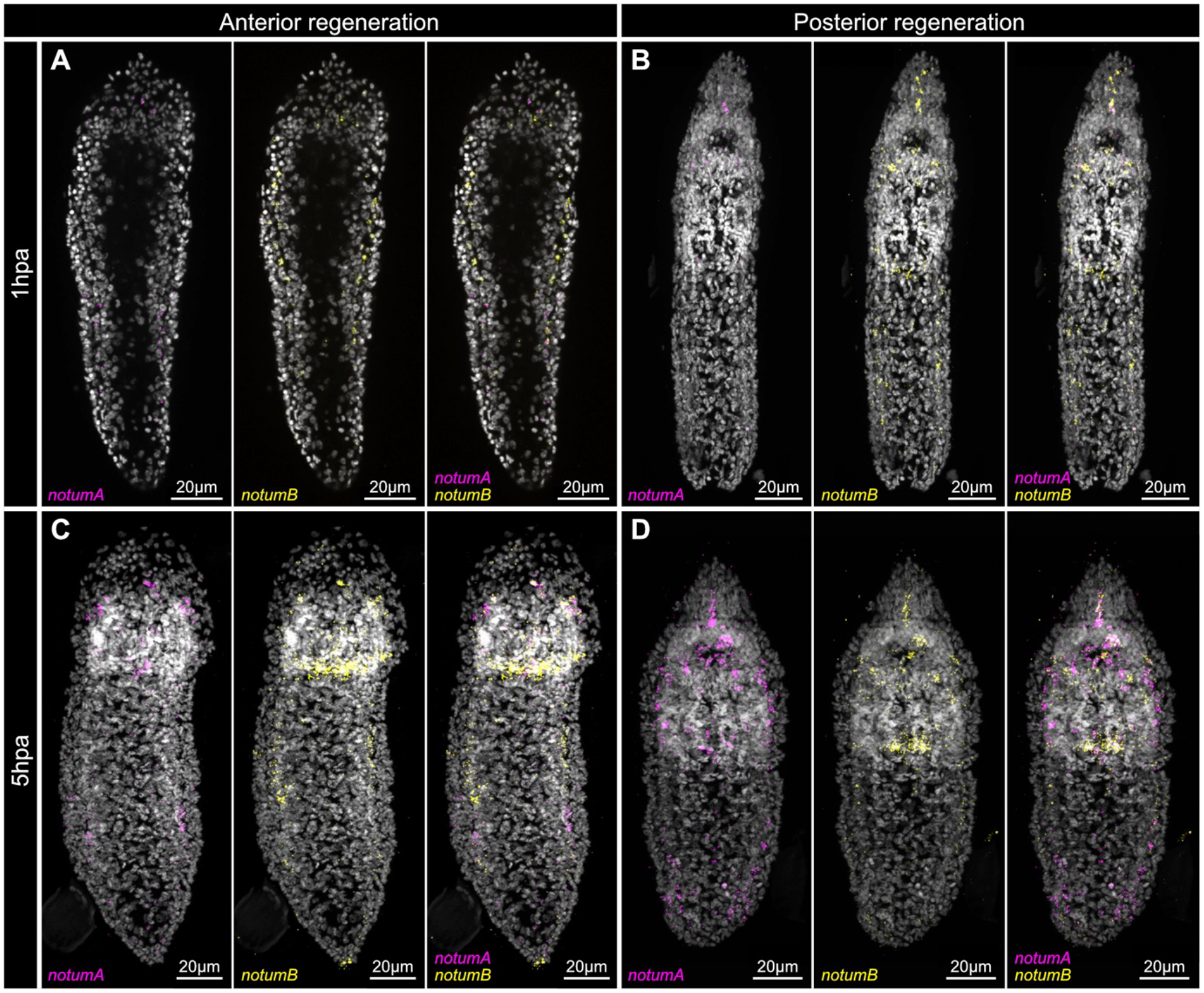
Expression of two paralogs of *notum* during head and tail regeneration of *Stenostomum brevipharyngium* at 1 and 5 hours post-amputation (hpa). Names of the hybridized genes are provided in the lower-left corner of each panel. Cell nuclei are counterstained with Hoechst (grey). Scale bars represent 20 μm.

### Function of Wnt signaling components in Stenostomum

Our recent research introduced dsRNA soaking as an efficient method for gene silencing in *S. brevipharyngium* (Gąsiorowski et al., 2025). To gain insight into the function of selected components of Wnt signaling in antero-posterior specification in *Stenostomum,* we performed RNAi of the posteriorly expressed Wnts (WntC1A, WntC1B), wound-activated Wnt (Wnt11), two β-catenin and two *notum* paralogs (Fig. 10). We first soaked the animals for 5 days in dsRNA targeted against each particular gene and then sorted the worms in three groups – intact, posterior regenerates and anterior regenerates – that were either soaked intact for additional 4 days or cut through the pharynx or posterior trunk, and then soaked throughout entire process of regeneration that takes up to 4 days.

**Fig. 10.**
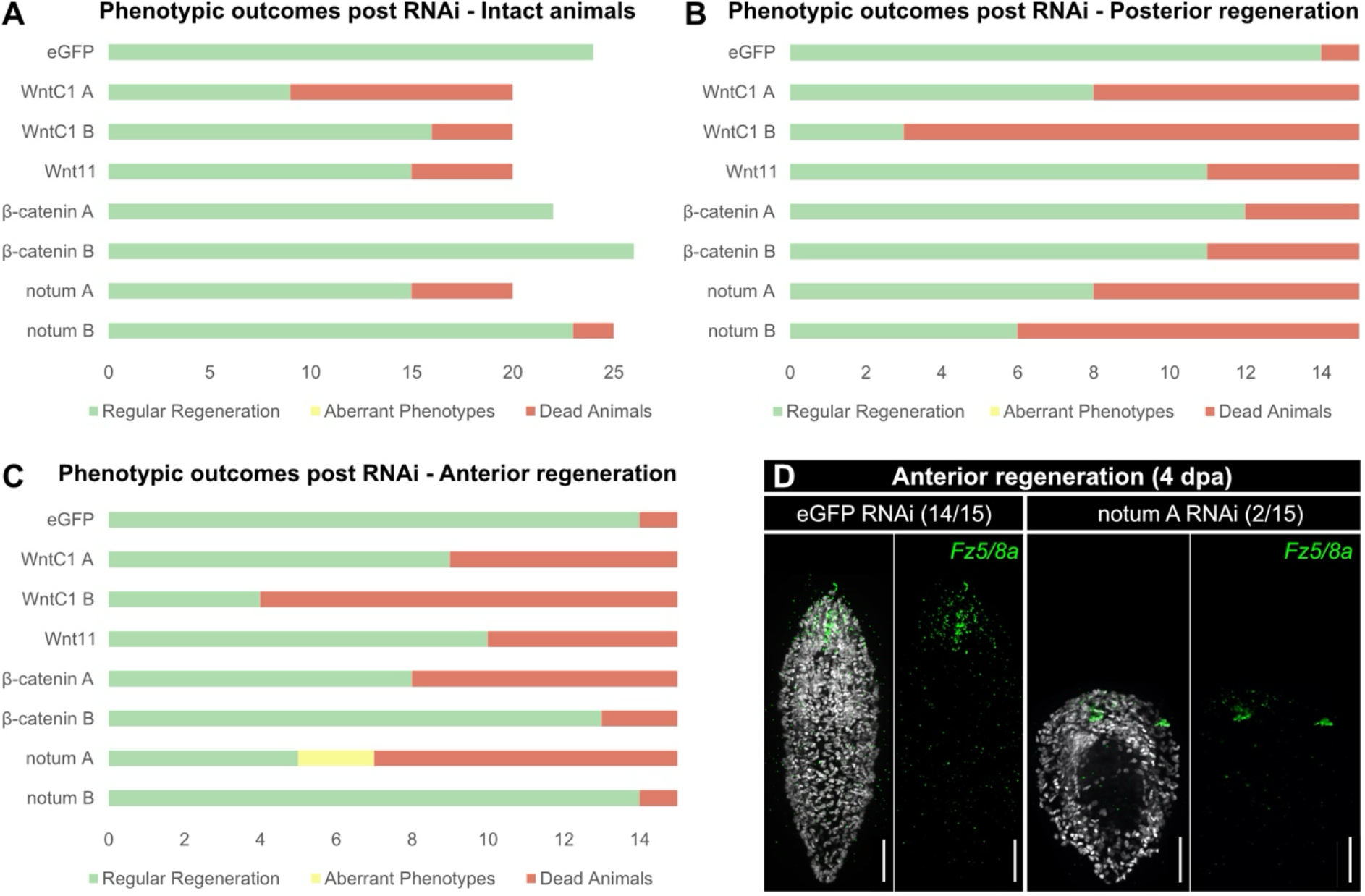
Functional analysis of the components of the Wnt signaling pathway using double-stranded RNA interference. Proportions of the phenotypic outcomes in intact animals (**A**), posterior regenerates (**B**), and anterior regenerates (**C**). **D.** Aberrant anterior regenerate in the *NotumA*(RNAi) compared to the control eGFP(RNAi) stained for cell nuclei with Hoechst (gray) and anterior marker Fz5/8a (green). Scale bars represent 20 μm.

In the intact group, most of the worms did not show phenotypes; however, some animals died in response to RNAi of Wnt and *notum* paralogs, with especially high mortality (60%) in the case of WntC1a (Fig. 10A). In the tail-regenerating worms, RNAi of all the genes resulted in moderate to high mortality, but we did not observe any overt phenotypesindicative of aberrant regeneration (Fig. 10B). Among head-regenerating worms, there was a relatively high degree of mortality in most of the RNAi experiments except *β-catenin B* and *notum B* (Fig. 10C). Additionally, two out of 15 worms in *notum A* (RNAi) showed aberrant head regeneration, resulting in anteriorly deformed worms (Fig. 10C and D). However, these worms still expressed the anterior marker *Fz5/8a* in their deformed anterior tissues (Fig. 10D), indicating that RNAi of *notum A* does not lead to posteriorization of the anterior wound and ectopic tail regeneration, as is the case in *S. mediterranea* (Petersen & Reddien, 2011). In general, RNAi of the components of Wnt signaling was either lethal or did not show any interpretable phenotypes. Importantly, our experiments did not indicate the reprogramming of tissue identity in response to experimental interference with canonical Wnt signaling, such as ectopic head induction in response to *β-catenin* knockdown or ectopic tail formation in response to *notum* knockdown. Whether this reflects differences in the specification of tissue identity during regeneration or technical shortcomings of the current RNAi protocol remains an open question.

## Discussion

Flatworms are famous for their regeneration abilities, including the ability to regenerate complete animals, with a fully functional head, from tiny tissue pieces (Egger et al., 2007; Ivankovic et al., 2019). However, while head regeneration has been studied extensively in regeneration-competent planarian species, the only other groups of flatworms known to include head regeneration-competent species are the distantly related microstomids and catenulids. This phylogenetic distribution raises an intriguing evolutionary question: Is head regeneration an ancestral trait in flatworms that was lost in most lineages or has head regeneration and the underlying molecular circuitry evolved convergently in only a few lineages? Addressing this question requires functional analysis of the key molecular circuits that drive regeneration in the different lineages (Srivastava, 2021).

While many aspects of flatworm regeneration remain unknown, the mechanistic analysis of head regeneration in the planarian model species over the last two decades has established Wnt signaling as a core molecular regulator of regeneration in planarians. High levels of Wnt signaling are necessary and sufficient for specifying tail identity, while the inhibition of Wnt signaling is necessary and sufficient for head specification. Wnt ligands and many of their regulators are expressed in molecular gradients along the A-P axis that are required for maintaining patterning and the anatomical identity along the axis (Almuedo-Castillo et al., 2012; Gurley et al., 2010; Petersen & Reddien, 2011; Stückemann et al., 2017). The gradients are thought to be organized by the poles, the anterior and posterior termini of the midline that express Wnt1 in the case of the tail pole and *notum* in the case of the head pole. Upon wounding or amputation, Wnt1 expression is prominently induced at all wounds irrespective of wound orientation (Almuedo-Castillo et al., 2012; Gurley et al., 2010; Petersen & Reddien, 2009), while the Wnt inhibitor *notum* is selectively expressed at anterior-facing wounds and functionally required as a symmetry-breaking factor (Petersen & Reddien, 2011). Many of the Wnt ligands and associated pathway components are expressed by the body wall musculature (Witchley et al., 2013), which surrounds all internal organs like a shell and is generally thought to act as a source of patterning information.

In light of the functional importance of Wnt signaling in planarians, the hallmarks of Wnt pathway deployment described above offer precise points of mechanistic comparison between regeneration-competent flatworm clades towards the goal of assessing conservation versus convergent evolution (Srivastava, 2021). The objective of this study was to generate a complementary data set of Wnt components and their expression pattern in our recently established *Stenostomum* laboratory model (Gąsiorowski et al., 2025), as a basis for future detailed mechanistic studies.

### Divergent evolution of Wnt and Frizzled-related proteins in flatworms

We first thought to systematically map the evolution of Wnt pathway components in flatworms and, specifically, the Wnt pathway complement in *Stenostomum*. By probing Wnt components in previously unstudied platyhelminth clades, we can now more precisely reconstruct the evolutionary history of their Wnt signaling system. For instance, we can confirm that six out of 13 ancestral bilaterian Wnt families (Wnt3, Wnt6, Wnt7, Wnt8, Wnt9 and Wnd10) were already lost before the divergence of contemporary Platyhelminthes. Such a loss of Wnt genes is not unusual, as a similar reduction has been reported in some other bilaterian clades, such as tardigrades (Chavarria et al., 2021) and insects (Janssen et al., 2010). Interestingly, catenulids retained WntA, which is missing in all the remaining flatworms (Rhabditophora), but have apparently lost Wnt1, which is otherwise well-conserved among rhabditophorans (Gurley et al., 2010; Riddiford & Olson, 2011).

Our extended sampling of Wnt ligands in multiple flatworm species further allowed classification of problematic planarian WntP-4 (also known as Wnt11-3 or Smed-Wnt-6) as an ortholog of Wnt 16, also present in catenulids but lacking in other Rhabditophora (Tab. 1). Altogether, it is evident that on top of the ancestral loss of six Wnt families in the common platyhelminth ancestor, particular clades of flatworms experienced independent losses of different Wnt orthologs. Despite these frequent losses, stenostomids and macrostomorphs have a high number of Wnt genes (encoding 11-14 proteins), comparable to other Metazoa with unreduced Wnt complements (Adamska et al., 2010; Borisenko et al., 2016; Cho et al., 2010; Holstein, 2012; Janssen et al., 2010; Kusserow et al., 2005; Lengfeld et al., 2009; Prud’homme et al., 2002; Pruitt et al., 2014; Riddiford & Olson, 2011; Robert et al., 2014; Somorjai et al., 2018; Vellutini et al., 2024), due to lineage-specific duplication and the presence of lineage-specific non-canonical Wnt ligands.

Frizzled-related proteins also show dynamic evolution within platyhelminths. Although all conserved families of Frizzled receptors are present in all analyzed flatworms, one of the canonical families of sFRPs (sFRP 3/4) has been lost in the last common platyhelminth ancestor. Both Frizzled receptors and sFRPs went through extensive expansions in flatworms, with multiple duplications of certain orthologs occurring independently in specific clades. These duplications resulted in a high number of both Frizzled receptors and sFRPs relative to ancestral bilaterian conditions (Adamska et al., 2010; Bastin et al., 2015; Robert et al., 2014; Schenkelaars et al., 2015; Vellutini et al., 2024).

Generally, the evolution of Wnt signaling components seems to have followed a pattern of ancestral loss of many conserved bilaterian gene families present in the last common platyhelminth ancestor, followed by additional lineage-specific losses and lineage-specific compensatory duplications of the remaining genes. A similar phenomenon has been observed for a number of other gene families in flatworms, such as Fox and Hox genes (Currie et al., 2016; Gąsiorowski, 2025; Gąsiorowski, Martín-Durán, et al., 2023; Pascual-Carreras et al., 2021). This is likely related to the ancestral reduction of the flatworm genome that resulted in massive losses of otherwise well-conserved genes (Grohme et al., 2018; Liao et al., 2023). Interestingly, in the case of Wnt signaling components, microscopic catenulids and macrostomorphs secondarily and independently evolved a complex complement of signaling molecules that is similar, at least by sheer gene number, to the ancestral bilaterian complement, and exceeds in this respect the systems present in large-bodied planarians and neodermatans.

### Similarities and differences in the expression of Wnt signaling components in flatworms

Most of the data on expression of Wnt signaling in various animal clades focuses on embryonic development (e.g., Adamska et al., 2010; Bastin et al., 2015; Cho et al., 2010; Janssen et al., 2010; Kusserow et al., 2005; Pruitt et al., 2014; Robert et al., 2014; Somorjai et al., 2018; Vellutini et al., 2024). However, in flatworms, Wnt signaling plays an important role in maintaining the antero-posterior axis during continuous development of tissue from adult pluripotent stem cells, and therefore, expression of pathway genes is well studied in adult animals (Armstrong et al., 2025; Gurley et al., 2010; Jarero et al., 2024; Stückemann et al., 2017). This allows direct comparison of the expression patterns of Wnt signaling components between *Stenostomum* and other platyhelminths.

Two of the Wnt5 paralogs in *Stenostomum* generally have an anterior expression domain, while in planarians, the gene is expressed on the edge of the body and in some neurons of the brain and nerve cords (Gurley et al., 2010). Interestingly, Wnt5 is also expressed in the lateral edges of the tapeworm larvae (Koziol et al., 2016), suggesting that this pattern might be a rhabditophoran innovation. A single paralog of Wnt11 in *Stenostomum* is expressed through the muscles of the trunk, showing, to some extent, a gradient towards the posterior tip, similarly to what has been reported for Wnt11 paralogs in planarians and tapeworms (Gurley et al., 2010; Jarero et al., 2024). Therefore, Wnt11 seems to have a conserved posterior expression among all tested groups of flatworms. As noted above, WntA, which is present in three copies in *Stenostomum*, has been lost in the rhabditophoran lineage, preventing direct comparison within platyhelminths. Similarly, there is no information on the expression of the Wnt16 ortholog in planarians, where it is known as Wnt11-3 or WntP-4, and the gene is missing from the other rhabditophorans tested thus far (Gurley et al., 2010; Riddiford & Olson, 2011).

It is also important to note here that three Wnt ligands in *Stenostomum* (Wnt16, WntC2b and WntC2a) show staggered expression in the trunk musculature that is repeated within the worm body. *Stenostomum* reproduces asexually through paratomy, a process in which new tail and head structures start to develop in the middle of the trunk, giving rise to two new individuals (da Rosa & Loreto, 2024; Gąsiorowski et al., 2025; Gąsiorowski, Dittmann, et al., 2023; Kepner & Cash, 1915; Ott, 1892; Sonneborn, 1930; Tratkiewicz & Gąsiorowski, 2025). Likely, these Wnt ligands play a role in establishing antero-posterior identities of the new individuals, and their staggered and repeated expression in the trunk may prepattern the asexual development of the zooids during paratomy.

Frizzled receptors in planarians have a wide range of expression patterns, with some of them showing antero-posterior regionalization, while others are expressed in the pharynx, nervous system, or throughout the entire body (Stückemann et al., 2017). Similarly, in tapeworms, Fz1/2/7 is expressed anteriorly, Fz4 posteriorly, and Fz5/8 has a complex expression that changes in different morphological sections of the animal (Jarero et al., 2024). In *Stenostomum,* in contrast, most Frizzled genes show staggered expression along the antero-posterior axis, with anterior restriction of three paralogs of Fz5/8, followed by the regions expressing Fz4 paralogs and expression of Fz9/10 in the trunk tissues, similarly to what has been proposed as the ancestral expression of these genes in bilaterian embryos (Vellutini et al., 2024). Therefore, it seems that despite the multiplication of some Frizzled genes in *Stenostomum*, their expression preserves ancestral bilaterian characteristics that have been lost in other flatworms.

sFRPs generally exhibit anterior expression in both planarians and tapeworms (Gurley et al., 2010; Gurley et al., 2008; Jarero et al., 2024), which is related to their putative function as antagonists of the posteriorly expressed Wnt ligands. However, sFRP1/2/5, which underwent extensive expansion in catenulids, shows very distinct expression for particular paralogs, with some of them marking the anterior structures, others active in the trunk, and a single paralog exclusively expressed in the tail. Even though sFRPs are believed to act primarily as Wnt antagonists, their function can be much more subtle, as they can also inhibit each other, bind to Frizzled receptors, and enhance diffusion of Wnt ligands (Bovolenta et al., 2008; Mii & Taira, 2009). We suggest that the expansion of sFRP1/2/5 in catenulids might be related to the subfunctionalization of particular paralogs and the emergence of a complex A/P patterning system that awaits further exploration.

Despite all the differences in expression of given components of Wnt signaling, a prevailing similarity between all flatworms is the expression of those genes in the musculature (Jarero et al., 2024; Witchley et al., 2013, this study). Information about expression of Wnt signaling genes in adult tissues is available only for a handful of invertebrate taxa (e.g., Borisenko et al., 2016; Lengfeld et al., 2009; Srivastava et al., 2014). Among these, Wnt signaling components are also expressed in muscles of acoels (Raz et al., 2017). Such similarity can either represent the ancient bilaterian condition conserved in both flatworms and acoelomorphs or an effect of convergent evolution. The latter might be related to similar developmental modes, as both clades rely on adult pluripotent stem cells for cellular turnover in their tissues (Gehrke & Srivastava, 2016), inherently dependent upon a system for positional patterning throughout adult life.

Altogether, even though Wnt ligands, their receptors, and regulators generally show antero-posterior expression gradients in the muscles of all flatworms, the deployment of particular orthologs into specific body regions seems to vary considerably between catenulids and rhabditophorans. This is not surprising, taking into account the long evolutionary distance between both clades and the complex evolutionary history of Wnt ligands in flatworms, marked by multiple lineage-specific losses and duplications that allowed for repetitive sub- and neofunctionalization of newly evolved paralogs.

### Conservation and divergence of wound-induced gene expression in flatworms

Immediate early response genes (IEGs), such as *Egr*, *Fos*, *Jun*, *Creb* and *Runt*, are activated at the wound site and initiate transcriptional responses to wounding (Srivastava, 2021). Interestingly, IEGs show remarkable conservation across metazoan phylogeny (Srivastava, 2021) with the set of homologous genes being activated in response to wounding in ctenophores (Mitchell et al., 2024), cnidarians (Cazet et al., 2021; Kaloulis et al., 2004), acoels (Gehrke et al., 2019), deuterostomes (Cary et al., 2020; Ishida et al., 2010; Martin & Nobes, 1992), and planarians (Wenemoser et al., 2012). In *Stenostomum*, multiple paralogs of *Egr*, *Fos*, *Jun* and *Runt* become upregulated during early stages of both anterior and posterior regeneration, and we documented elevated expression of *Runt-1A* at wound sites 1 hpa. Therefore, the ancestral role of IEGs in early wound response seems to be well conserved in catenulids.

In planarians, IEG activation is followed by expression of Wnt1 at any wound site (Almuedo-Castillo et al., 2012; Gurley et al., 2010; Petersen & Reddien, 2008). In *Stenostomum*, we also observed a single Wnt ligand activated upon both anterior and posterior wounding; however, it was not an ortholog of Wnt1, which is generally missing in catenulids, but a catenulid-specific Wnt protein – WntC1a. This indicates that despite general mechanistic similarity between *Stenostomum* and *Schmidtea*, deployment of particular Wnt ligands is evolutionarily labile, suggesting possible developmental system drift (True & Haag, 2001) or convergent, independent evolution of wound-induced Wnt expression. Moreover, expression of WntC1a in *Stenostomum* is not restricted to the wound site, as is the case for Wnt1 in planarians. Instead, WntC1a shows broader expression upon injury that spreads throughout most of the animal. We suggest that differences in body size can explain this dissimilarity. *S. brevipharyngium* is a microscopic animal, with a body length of ca. 150-250 μm. Therefore, its entire body is inevitably proximal to the wound when compared to planarians that are roughly a hundred times longer (15-20 mm). Wound-induced WntC1a expression is restricted to the musculature in *Stenostomum,* and several of its longitudinal muscle fibers span the entire trunk of the worm. Therefore, wounding of one such long muscle fiber – regardless of the wound site – could evoke transcriptional responses in the entire length of the animal. This contrast between microscopic catenulids and macroscopic planarians highlights that body size should be taken into account when comparing cellular mechanisms of regeneration between organisms.

One of the hallmarks of planarian regeneration is the *notum*-mediated inhibition of *β-catenin/*Wnt signaling at the anterior wound site (Petersen & Reddien, 2011). At the posterior wound, where β-catenin remains active, paralogs of the posterior Wnt11 are expressed, leading to the development of tail tissues (Almuedo-Castillo et al., 2012; Petersen & Reddien, 2008). Our results suggest divergent regulation of these genes in *Stenostomum*. Even though two *notum* paralogs are expressed in the anterior tissues of intact animals, they do not show similar dynamics during regeneration. Instead, both genes show weaker expression at the early stage of regeneration (1 hpa) and are expressed at 5 hpa in both anteriorly and posteriorly regenerating animals. Wnt11 is expressed in the posterior trunk of intact *Stenostomum*, similar to its expression in intact planarians, but it is upregulated during both head and tail regeneration in *Stenostomum*. Finally, our RNAi experiments, in which we knock down expression of *notum*, *β-catenin* and posterior Wnts, did not recreate double-head or double-tail phenotypes typical for knock-downs of those genes in regenerating planarians (Almuedo-Castillo et al., 2012; Gurley et al., 2008; Iglesias et al., 2008; Petersen & Reddien, 2008), even though such phenotypes are generally known to occur in *Stenostomum* (Sonneborn, 1930; Tratkiewicz & Gąsiorowski, 2025). This could be either related to a fundamentally different regulatory logic of the head regeneration process between planarians and catenulids or to the fact that these genes are present in multiple copies in *S. brevipharyngium*, which allows for functional compensation between paralogs.

Altogether, our data indicate that while Wnt signaling is involved in the regeneration process in catenulids, it differs remarkably from the system known in planarians. In both clades, there is a wound-responsive Wnt ligand, but it is not homologous between the two groups, and the Notum/β-catenin/Wnt regulatory network does not seem to be directly involved in the specification of anterior vs posterior fate during catenulid regeneration. Interestingly, a recent survey of β-catenin in another flatworm, *Macrostomum*, also indicated that the role of this gene in the regeneration process is different from that in planarians (Kibet et al., 2025). Therefore, it seems that the exact involvement of Wnt signaling in the regeneration process is evolutionarily labile within flatworms, and its strict and crucial role in the specification of anterior vs posterior fate might be restricted to planarians.

## Conclusions

Wnt signaling underwent extensive and largely divergent evolution within flatworms, which is evident in the complement of pathway components, the spatial patterns of their expression in intact animals, and their temporal dynamics and functions during the regeneration process. Such dissimilarity hints at independent evolution of head regeneration between catenulids and planarians, in accordance with the phylogenetic distribution of head regenerative capacities in flatworms. Alternatively, observed differences in the expression dynamics and functions of particular Wnt signaling components between both clades might be explained by differences in body size or developmental system drift driven by independent evolutionary pressures that modified the ancestral homologous regulatory network beyond recognition. Either way, the dissimilarities resulting from the divergent evolution of the platyhelminth Wnt system restrict the usefulness of direct comparison of Wnt function in the inference of homology of regeneration processes in flatworms.

## Materials and Methods

### Search for orthologs and phylogenetic analyses

We performed a reciprocal BLAST search for sequences encoding putative Wnt, Frizzled and sFRP proteins in the published transcriptomes of two catenulids, *S. brevipharyngium* (PRJNA1004231 published in Gąsiorowski, Dittmann, et al. (2023)) and *S. lecuops* (PRJNA276469 published in Laumer et al. (2015)), as well as genomes of two species of *Macrostomum*, *M. cliftonense* and *M. hystrix* published in Wiberg et al. (2023). We also searched for Frizzled and sFRP genes in the transcriptomes of *S. mediterranea* and four neodermatan species available in the Plan Mine database (Brandl et al., 2016). To complement our dataset on Wnt signaling components in flatworms, we also use published Wnt sequences from *S. mediterranea* and neodermatans (Riddiford & Olson, 2011). Gene IDs of all newly identified genes are available in Table S1.

To assign the obtained sequences into respective conserved metazoan families, we aligned them with metazoan reference sequences (Tables S2 and S3) using Clustal Omega (v 1.2.3) (Sievers & Higgins, 2014) implemented in Geneious Prime (v2023.0.3), creating one alignment for all Wnt genes and another for Frizzled receptors and related proteins (sFRPs and smoothened). The alignments were then trimmed in Geneious Prime to remove sites containing more than 50% gaps (trimmed alignments are available as Supplementary Data 1 and 2) and analyzed with raxmlGUI (v 2.0) (Edler et al., 2021) to generate phylogenetic trees. The LG amino acid substitution model was used for both analyses based on the results of ModelTest-NG v0.1.7 (Darriba et al., 2020) implemented in raxmlGUI. Both analyses were run with 100 bootstrap replicates. Phylogenetic trees with terminal names and all bootstrap support values are provided as Figures S2 and S3.

### Animal husbandry

*S. brevipharyngium* cultures were ordered in 2010 from Connecticut Valley Biological Supply and have been maintained in the laboratory since then. The cultures were maintained in standardized Chalkley’s Medium (CM) at 20°C in the dark and fed *ad libitum* with the unicellular eukaryote *Chilomonas paramecium*.

### HCR in situ hybridization

Animals were first anesthetized in 1.44% (w:v) MgCl_2_ in CM for approximately 10 min until they stopped active movements. Then, the worms were fixed in 4% (v:v) formaldehyde in PBS + 0.1% Tween-20 detergent (PTw) for 30 minutes, washed three times in PTw, dehydrated in 100% methanol and stored in fresh 100% methanol in −20°C for at least 12h.

The DNA probe oligo pools for fluorescent RNA in situ hybridization chain reaction (HCR) v3.0 (Choi et al., 2018) were designed using the Özpolat Lab’s in situ probe generator (Kuehn et al., 2022) and ordered from Integrated DNA Technologies. For in situ staining, the worms were rehydrated in Me-OH/PTw series at RT, washed four times in PTw, and then prehybridized in hybridization buffer for 40 min at 37oC. Next, the animals were placed in HCR probe mixtures at 1uM concentration in hybridization buffer and incubated over night at 37°C. On the next day, the probes were removed with four washes of probe wash solution (each 10 min at 37°C), followed by three washes in 5xSSC+0.1% Tween-20 at RT. Then, the worms were incubated in amplification buffer for 30 min at RT. In the meantime, the HCR hairpins (ordered from Molecular Instruments) were prepared by heating for 1min 30sec at 95°C and then cooled down at RT in the dark for 30 min. Animals were incubated in the amplification buffer with hairpins at 40nM concentration overnight at RT in the dark. On the next day, the samples were rinsed three times with 5xSSC+0.1% Tween-20, twice with PTw and incubated for 40 min in Hoechst 33342 in PTw (1:5000). Stained specimens were washed once in PBS, mounted in Fluoromount G (Thermo Fischer, 00-4958-02), left overnight at 4°C and then imaged with an Olympus IX83 microscope with a spinning disc Yokogawa CSUW1-T2S scan head.

### Amputations and regeneration

For amputation, worms were anesthetized with 1% (w:v) MgCl_2_ hexahydrate in CM for ca. 10 min., cut under a dissecting scope with an eyelash at the level of the pharynx (anterior regeneration) or through the posterior half of the trunk (posterior regeneration), and immediately transferred to fresh CM. The regenerates were washed a few times with CM to remove the anesthetic medium and then kept in the dark at 20 °C until processing.

### RNA sequencing and analysis

For each RNA extraction, about 60 worms were pooled and preserved in RNA-later. RNA extraction was performed using the NucleoSpin RNA XS kit for RNA purification, Macherey–Nagel, 740902.50, according to the manufacturer’s recommendation, with the following modifications: before extraction, RNAase-free PBS was added to each sample to dilute the RNA-later reagent, and the samples were centrifuged at 4500 × G for 10 min at 4 °C in a benchtop centrifuge to pellet the worm tissues. Then the supernatant was removed and replaced with the kit’s lysis buffer. To enhance the lysis, samples were snap-frozen in liquid nitrogen and then thawed in a water bath at room temperature. Next, the samples were processed following the standard protocol for RNA isolation from tissues. The extracted RNA was evaluated for concentration and integrity on an Agilent 2100 Bioanalyzer with an RNA 6000 Nano chip and sent for sequencing on an Illumina NovaSeq 6000 at the Dresden Concept Genome Center to a depth of 40 million paired-end reads with 100 bp length.

Quality control of sequencing data was performed using FastQC (v0.11.9) (Andrews, 2010) before and after trimming. Trimming was conducted using the Trimmomatic tool (v0.39) (Bolger et al., 2014). A contamination screen was performed with Kraken2 (v2.1.2) (Wood et al., 2019), using the reference database k2_standard_20201202.tar.gz (https://genome-idx.s3.amazonaws.com/kraken/k2_standard_20201202.tar.gz).

Transcriptome alignment was carried out using the Trinity suite (v2.14.0) (Grabherr et al., 2011), with the parameters ‘--est_method RSEM’ and ‘--aln_method bowtie2’. For alignment, the assembled reference transcriptome of *S. brevipharyngium* (go_Sbre_v1) (Gąsiorowski, Dittmann, et al., 2023) was used, available via Zenodo (DOI: 10.5281/zenodo.8239273).

Transcripts with low or no expression were filtered prior to downstream analysis. Specifically, only transcripts with at least five reads across a minimum of three samples were retained. Read counts were normalized based on library size (Chen et al., 2025) and subsequently subjected to regularized log transformation (Love et al., 2014) for PCA visualization (Lê et al., 2008). Differential expression analysis was performed using the Limma package (Ritchie et al., 2015), applying cut-off thresholds of an adjusted p-value < 0.05 and an absolute value of log₂ fold change (L2FC) ≥ 1. Lists of genes differentially expressed for each comparison, with automatic annotation, logFC, and adjusted P-values are provided in Tables S6-S15.

### Data deposition

The raw RNA-seq reads of regenerating worms have been deposited at the NCBI Sequence Reads Archive as BioProject PRJNA1492546.

### RNAi

Target gene sequences were amplified from *S. brevipharyngium* cDNA via PCR and inserted into the pRTP4P-2.0 vector using Gibson assembly. The recombinant plasmids were introduced into *E. coli* DH5α cells via heat-shock transformation. The transformed colonies were selected, and plasmid DNA was extracted using the standard NucleoSpin miniprep kit (MN-740588.250). Constructs were confirmed by sequencing and used as templates for PCR with PR244 and T7AA18 primers and Q5 High-Fidelity DNA Polymerase (NEB, M0491S). To reach the desired concentration, two rounds of PCR reactions with 35 cycles each were required. The specificity of the reaction was checked with DNA gel electrophoresis.

The purified PCR products were used as templates to produce dsRNA with T7 RNA polymerase (Thermo Fisher, EP0112) in an in-vitro RNA transcription reaction. The synthesized RNA was annealed by heating to 75°C and then cooling down to room temperature. The concentration of the dsRNA was measured on a NanoDrop ND-1000, and its integrity was checked with gel electrophoresis.

Administration of dsRNA followed a previously established protocol (Gąsiorowski et al., 2025). The dsRNA obtained was diluted in 600μL of Chalkey’s Medium (CM), with *Chilomonas paramecium* solution as a food source, to a concentration of 100 ng/μL and distributed into 24 well plates. Approximately 50 worms were put in each well. Following exposure to dsRNA for 5 days, the worms were amputated and left to regenerate for 4 days, after which the effects of the dsRNA were studied using a dissecting microscope and HCR in situ hybridization. Every day for the duration of the experiment, 400μL of the dsRNA solution was replaced in the wells, with additional food added when *Chilomonas* were no longer present. The integrity of the freshly added dsRNA was regularly checked with gel electrophoresis.

## Acknowledgements

All current and past members of Rink Laboratory are acknowledged for their scientific and technical support. Especially, we would like to thank Jason Pellettieri for his help with editing the draft of the manuscript. The research was supported by the Alexander von Humboldt Foundation (The Humboldt Research Fellowship for Postdoctoral Researchers to L.G.) and the Max Planck Society (funding to J.C.R.).

## Notes

### Competing Interest Statement

The authors have declared no competing interest.

## References

Adamska, M., Larroux, C., Adamski, M., Green, K., Lovas, E., Koop, D., Richards, G. S., Zwafink, C., & Degnan, B. M. (2010). Structure and expression of conserved Wnt pathway components in the demosponge Amphimedon queenslandica. Evolution & Development, 12(5), 494–518. 10.1111/j.1525-142X.2010.00435.x

Almuedo-Castillo, M., Sureda-Gómez, M., & Adell, T. (2012). Wnt signaling in planarians: new answers to old questions. International Journal of Developmental Biology, 56(1-2-3), 53–65.

Andrews, S. (2010). Fast QC: a quality control tool for high throughput sequence data. Retrieved 15.01.2023 from http://www.bioinformatics.babraham.ac.uk/projects/fastqc/

Armstrong, R., Marks, N. J., Geary, T. G., Harrington, J., Selzer, P. M., & Maule, A. G. (2025). Wnt/β-catenin signalling underpins juvenile Fasciola hepatica growth and development. PLoS pathogens, 21(2), e1012562.

Bastin, B. R., Chou, H.-C., Pruitt, M. M., & Schneider, S. Q. (2015). Structure, phylogeny, and expression of the frizzled-related gene family in the lophotrochozoan annelid Platynereis dumerilii. EvoDevo, 6(1), 37. 10.1186/s13227-015-0032-4

Bolger, A. M., Lohse, M., & Usadel, B. (2014). Trimmomatic: a flexible trimmer for Illumina sequence data. Bioinformatics, 30(15), 2114–2120.

Borisenko, I., Adamski, M., Ereskovsky, A., & Adamska, M. (2016). Surprisingly rich repertoire of Wnt genes in the demosponge Halisarca dujardini. BMC Evolutionary Biology, 16(1), 123. 10.1186/s12862-016-0700-6

Bovolenta, P., Esteve, P., Ruiz, J. M., Cisneros, E., & Lopez-Rios, J. (2008). Beyond Wnt inhibition: new functions of secreted Frizzled-related proteins in development and disease. Journal of cell science, 121(6), 737–746. 10.1242/jcs.026096

Brandl, H., Moon, H., Vila-Farré, M., Liu, S.-Y., Henry, I., & Rink, J. C. (2016). PlanMine–a mineable resource of planarian biology and biodiversity. Nucleic acids research, 44(D1), D764–D773.

Cary, G. A., McCauley, B. S., Zueva, O., Pattinato, J., Longabaugh, W., & Hinman, V. F. (2020). Systematic comparison of sea urchin and sea star developmental gene regulatory networks explains how novelty is incorporated in early development. Nature Communications, 11(1), 6235.

Cazet, J. F., Cho, A., & Juliano, C. E. (2021). Generic injuries are sufficient to induce ectopic Wnt organizers in Hydra. Elife, 10, e60562. 10.7554/eLife.60562

Chavarria, R. A., Game, M., Arbelaez, B., Ramnarine, C., Snow, Z. K., & Smith, F. W. (2021). Extensive loss of Wnt genes in Tardigrada. BMC ecology and evolution, 21(1), 223.

Chen, Y., Chen, L., Lun, A. T., Baldoni, P. L., & Smyth, G. K. (2025). edgeR v4: powerful differential analysis of sequencing data with expanded functionality and improved support for small counts and larger datasets. Nucleic acids research, 53(2), gkaf018.

Cho, S.-J., Vallès, Y., Giani, V. C., Jr, Seaver, E. C., & Weisblat, D. A. (2010). Evolutionary Dynamics of the wnt Gene Family: A Lophotrochozoan Perspective. Molecular biology and evolution, 27(7), 1645–1658. 10.1093/molbev/msq052

Choi, H. M. T., Schwarzkopf, M., Fornace, M. E., Acharya, A., Artavanis, G., Stegmaier, J., Cunha, A., & Pierce, N. A. (2018). Third-generation in situ hybridization chain reaction: multiplexed, quantitative, sensitive, versatile, robust. Development, 145(12). 10.1242/dev.165753

Currie, K. W., Brown, D. D., Zhu, S., Xu, C., Voisin, V., Bader, G. D., & Pearson, B. J. (2016). HOX gene complement and expression in the planarian *Schmidtea mediterranea*. EvoDevo, 7, 7. 10.1186/s13227-016-0044-8

da Rosa, M. T., & Loreto, E. L. (2022). Revisiting the regeneration of *Stenostomum leucops* (Catenulida, Platyhelminthes). Invertebrate Reproduction & Development, 66(1), 1–7.

da Rosa, M. T., & Loreto, E. L. S. (2024). Mining differentially expressed genes during paratomy in the transcriptome of the flatworm *Stenostomum leucops*. *Scientific reports*, *14*(1), 29267.

Darriba, D., Posada, D., Kozlov, A. M., Stamatakis, A., Morel, B., & Flouri, T. (2020). ModelTest-NG: a new and scalable tool for the selection of DNA and protein evolutionary models. Molecular biology and evolution, 37(1), 291–294.

Dirks, U., Gruber-Vodicka, H. R., Egger, B., & Ott, J. A. (2012). Proliferation pattern during rostrum regeneration of the symbiotic flatworm *Paracatenula galateia*: a pulse-chase-pulse analysis. Cell and Tissue Research, 349(2), 517–525. 10.1007/s00441-012-1426-4

Edler, D., Klein, J., Antonelli, A., & Silvestro, D. (2021). raxmlGUI 2.0: A graphical interface and toolkit for phylogenetic analyses using RAxML. Methods in Ecology and Evolution, 12(2), 373–377. 10.1111/2041-210X.13512

Egger, B., Gschwentner, R., & Rieger, R. (2007). Free-living flatworms under the knife: past and present. Development genes and evolution, 217, 89–104.

Egger, B., Lapraz, F., Tomiczek, B., Müller, S., Dessimoz, C., Girstmair, J., Škunca, N., Rawlinson, K. A., Cameron, C. B., & Beli, E. (2015). A transcriptomic-phylogenomic analysis of the evolutionary relationships of flatworms. Current Biology, 25(10), 1347–1353.

Gąsiorowski, L. (2025). Evidence for multiple independent expansions of Fox gene families within flatworms. Journal of Molecular Evolution, 1–12.

Gąsiorowski, L., Chai, C., Rozanski, A., Purandare, G., Ficze, F., Mizi, A., Wang, B., & Rink, J. C. (2025). Regeneration in the absence of canonical neoblasts in an early branching flatworm. Nature Communications, 16(1), 1232.

Gąsiorowski, L., Dittmann, I. L., Brand, J. N., Ruhwedel, T., Möbius, W., Egger, B., & Rink, J. C. (2023). Convergent evolution of the sensory pits in and within flatworms. BMC biology, 21(1), 1–19.

Gąsiorowski, L., Martín-Durán, J. M., & Hejnol, A. (2023). The evolution of Hox genes in Spiralia. In D. E. K. Ferrier (Ed.), Hox Modules in Evolution and Development (pp. 177– 194). CRC Press.

Gehrke, A. R., Neverett, E., Luo, Y.-J., Brandt, A., Ricci, L., Hulett, R. E., Gompers, A., Ruby, J. G., Rokhsar, D. S., Reddien, P. W., & Srivastava, M. (2019). Acoel genome reveals the regulatory landscape of whole-body regeneration. Science, 363(6432), eaau6173. doi:10.1126/science.aau6173

Gehrke, A. R., & Srivastava, M. (2016). Neoblasts and the evolution of whole-body regeneration. Current opinion in genetics & development, 40, 131–137.

Grabherr, M. G., Haas, B. J., Yassour, M., Levin, J. Z., Thompson, D. A., Amit, I., Adiconis, X., Fan, L., Raychowdhury, R., & Zeng, Q. (2011). Full-length transcriptome assembly from RNA-Seq data without a reference genome. Nature biotechnology, 29(7), 644–652.

Grohme, M. A., Schloissnig, S., Rozanski, A., Pippel, M., Young, G. R., Winkler, S., Brandl, H., Henry, I., Dahl, A., Powell, S., Hiller, M., Myers, E., & Rink, J. C. (2018). The genome of Schmidtea mediterranea and the evolution of core cellular mechanisms. Nature, 554(7690), 56–61. 10.1038/nature25473

Gurley, K. A., Elliott, S. A., Simakov, O., Schmidt, H. A., Holstein, T. W., & Alvarado, A. S. (2010). Expression of secreted Wnt pathway components reveals unexpected complexity of the planarian amputation response. Developmental biology, 347(1), 24–39.

Gurley, K. A., Rink, J. C., & Alvarado, A. S. (2008). β-catenin defines head versus tail identity during planarian regeneration and homeostasis. Science, 319(5861), 323–327.

Holstein, T. W. (2012). The evolution of the Wnt pathway. Cold Spring Harbor Perspectives in Biology, 4(7), a007922.

Iglesias, M., Gomez-Skarmeta, J. L., Saló, E., & Adell, T. (2008). Silencing of Smed-β catenin1 generates radial-like hypercephalized planarians. Development, 135(7), 1215–1221.

Ishida, T., Nakajima, T., Kudo, A., & Kawakami, A. (2010). Phosphorylation of Junb family proteins by the Jun N-terminal kinase supports tissue regeneration in zebrafish. Developmental biology, 340(2), 468–479. 10.1016/j.ydbio.2010.01.036

Ivankovic, M., Haneckova, R., Thommen, A., Grohme, M. A., Vila-Farré, M., Werner, S., & Rink, J. C. (2019). Model systems for regeneration: planarians. Development, 146(17). 10.1242/dev.167684

Janssen, R., Le Gouar, M., Pechmann, M., Poulin, F., Bolognesi, R., Schwager, E. E., Hopfen, C., Colbourne, J. K., Budd, G. E., Brown, S. J., Prpic, N.-M., Kosiol, C., Vervoort, M., Damen, W. G. M., Balavoine, G., & McGregor, A. P. (2010). Conservation, loss, and redeployment of Wnt ligands in protostomes: implications for understanding the evolution of segment formation. BMC Evolutionary Biology, 10(1), 374. 10.1186/1471-2148-10-374

Jarero, F., Baillie, A., Riddiford, N., Montagne, J., Koziol, U., & Olson, P. D. (2024). Muscular remodeling and anteroposterior patterning during tapeworm segmentation. Developmental Dynamics, 253(11), 998–1023. 10.1002/dvdy.712

Kaloulis, K., Chera, S., Hassel, M., Gauchat, D., & Galliot, B. (2004). Reactivation of developmental programs: the cAMP-response element-binding protein pathway is involved in hydra head regeneration. Proceedings of the National Academy of Sciences, 101(8), 2363–2368.

Kepner, W. A., & Cash, J. (1915). Ciliated pits of *Stenostoma*. Journal of Morphology, 26(2), 235–245.

Kibet, M. K., Hilchenbach, J., Neumann, L., Mayer, R., Aigner, G. P., Höckner, M., Hobmayer, B., & Egger, B. (2025). Silencing of β-catenin1 blocks tail regeneration, but does not induce head regeneration in the flatworm *Macrostomum lignano*. Discover Developmental Biology, 235(1), 1–21.

Koziol, U., Jarero, F., Olson, P. D., & Brehm, K. (2016). Comparative analysis of Wnt expression identifies a highly conserved developmental transition in flatworms. BMC biology, 14(1), 10. 10.1186/s12915-016-0233-x

Kuehn, E., Clausen, D. S., Null, R. W., Metzger, B. M., Willis, A. D., & Özpolat, B. D. (2022). Segment number threshold determines juvenile onset of germline cluster expansion in *Platynereis dumerilii*. Journal of Experimental Zoology Part B: Molecular and Developmental Evolution, 338(4), 225–240.

Kusserow, A., Pang, K., Sturm, C., Hrouda, M., Lentfer, J., Schmidt, H. A., Technau, U., von Haeseler, A., Hobmayer, B., Martindale, M. Q., & Holstein, T. W. (2005). Unexpected complexity of the Wnt gene family in a sea anemone. Nature, 433(7022), 156–160. 10.1038/nature03158

Laumer, C. E., Hejnol, A., & Giribet, G. (2015). Nuclear genomic signals of the ’microturbellarian’ roots of platyhelminth evolutionary innovation. Elife, 4. 10.7554/eLife.05503

Lê, S., Josse, J., & Husson, F. (2008). FactoMineR: an R package for multivariate analysis. Journal of statistical software, 25, 1–18.

Lengfeld, T., Watanabe, H., Simakov, O., Lindgens, D., Gee, L., Law, L., Schmidt, H. A., Özbek, S., Bode, H., & Holstein, T. W. (2009). Multiple Wnts are involved in Hydra organizer formation and regeneration. Developmental biology, 330(1), 186–199. 10.1016/j.ydbio.2009.02.004

Liao, I. J.-Y., Lu, T.-M., Chen, M.-E., & Luo, Y.-J. (2023). Spiralian genomics and the evolution of animal genome architecture. Briefings in Functional Genomics, 22(6), 498–508.

Liu, S.-Y., Selck, C., Friedrich, B., Lutz, R., Vila-Farré, M., Dahl, A., Brandl, H., Lakshmanaperumal, N., Henry, I., & Rink, J. C. (2013). Reactivating head regrowth in a regeneration-deficient planarian species. Nature, 500(7460), 81–84.

Love, M. I., Huber, W., & Anders, S. (2014). Moderated estimation of fold change and dispersion for RNA-seq data with DESeq2. Genome Biology, 15(12), 550.

Martin, P., & Nobes, C. D. (1992). An early molecular component of the wound healing response in rat embryos—induction of c-fos protein in cells at the epidermal wound margin. Mechanisms of development, 38(3), 209–215.

Mii, Y., & Taira, M. (2009). Secreted Frizzled-related proteins enhance the diffusion of Wnt ligands and expand their signalling range. Development, 136(24), 4083–4088. 10.1242/dev.032524

Mitchell, D. G., Edgar, A., Mateu, J. R., Ryan, J. F., & Martindale, M. Q. (2024). The ctenophore *Mnemiopsis leidyi* deploys a rapid injury response dating back to the last common animal ancestor. Communications Biology, 7(1), 203.

Moraczewski, J. (1977). Asexual reproduction and regeneration of *Catenula* (Turbellaria, Archoophora). Zoomorphologie, 88(1), 65–80.

Newmark, P. A., & Alvarado, A. S. (2002). Not your father’s planarian: a classic model enters the era of functional genomics. Nature Reviews Genetics, 3(3), 210–219. 10.1038/nrg759

Ott, H. N. (1892). A study of *Stenostoma leucops* O. Schm. Journal of Morphology, 7(3), 263–304. 10.1002/jmor.1050070302

Palmberg, I. (1986). Cell migration and differentiation during wound healing and regeneration in *Microstomum lineare* (Turbellaria). Advances in the Biology of Turbellarians and Related Platyhelminthes: Proceedings of the Fourth International Symposium on the Turbellaria held at Fredericton, New Brunswick, Canada, August 5–10, 1984,

Pascual-Carreras, E., Herrera-Úbeda, C., Rosselló, M., Coronel-Córdoba, P., Garcia-Fernàndez, J., Saló, E., & Adell, T. (2021). Analysis of Fox genes in *Schmidtea mediterranea* reveals new families and a conserved role of Smed-foxO in controlling cell death. Scientific reports, 11(1), 2947.

Petersen, C. P., & Reddien, P. W. (2008). Smed-β catenin-1 is required for anteroposterior blastema polarity in planarian regeneration. Science, 319(5861), 327–330.

Petersen, C. P., & Reddien, P. W. (2009). A wound-induced Wnt expression program controls planarian regeneration polarity. Proceedings of the National Academy of Sciences, 106(40), 17061–17066.

Petersen, C. P., & Reddien, P. W. (2011). Polarized notum activation at wounds inhibits Wnt function to promote planarian head regeneration. Science, 332(6031), 852–855.

Prud’homme, B., Lartillot, N., Balavoine, G., Adoutte, A., & Vervoort, M. (2002). Phylogenetic Analysis of the Wnt Gene Family: Insights from Lophotrochozoan Members. Current Biology, 12(16), 1395–1400. 10.1016/S0960-9822(02)01068-0

Pruitt, M. M., Letcher, E. J., Chou, H.-C., Bastin, B. R., & Schneider, S. Q. (2014). Expression of the wnt gene complement in a spiral-cleaving embryo and trochophore larva. Int J Dev Biol, 58(6-8), 563–573.

Raz, A. A., Srivastava, M., Salvamoser, R., & Reddien, P. W. (2017). Acoel regeneration mechanisms indicate an ancient role for muscle in regenerative patterning. Nature Communications, 8(1), 1260. 10.1038/s41467-017-01148-5

Riddiford, N., & Olson, P. D. (2011). Wnt gene loss in flatworms. Development genes and evolution, 221, 187–197.

Ritchie, M. E., Phipson, B., Wu, D., Hu, Y., Law, C. W., Shi, W., & Smyth, G. K. (2015). limma powers differential expression analyses for RNA-sequencing and microarray studies. Nucleic acids research, 43(7), e47–e47.

Robert, N., Lhomond, G., Schubert, M., & Croce, J. C. (2014). A comprehensive survey of wnt and frizzled expression in the sea urchin *Paracentrotus lividus*. genesis, 52(3), 235–250. 10.1002/dvg.22754

Schenkelaars, Q., Fierro-Constain, L., Renard, E., Hill, A. L., & Borchiellini, C. (2015). Insights into Frizzled evolution and new perspectives. Evol Dev, 17(2), 160–169. 10.1111/ede.12115

Sievers, F., & Higgins, D. G. (2014). Clustal Omega, Accurate Alignment of Very Large Numbers of Sequences. In D. J. Russell (Ed.), Multiple Sequence Alignment Methods (pp. 105–116). Humana Press. 10.1007/978-1-62703-646-7_6

Sikes, J. M., & Newmark, P. A. (2013). Restoration of anterior regeneration in a planarian with limited regenerative ability. Nature, 500(7460), 77–80. 10.1038/nature12403

Somorjai, I. M. L., Martí-Solans, J., Diaz-Gracia, M., Nishida, H., Imai, K. S., Escrivà, H., Cañestro, C., & Albalat, R. (2018). Wnt evolution and function shuffling in liberal and conservative chordate genomes. Genome Biology, 19(1), 98. 10.1186/s13059-018-1468-3

Sonneborn, T. M. (1930). Genetic studies on *Stenostomum incaudatum* (nov. spec.). I. The nature and origin of differences among individuals formed during vegetative reproduction. Journal of Experimental Zoology, 57(1), 57–108.

Srivastava, M. (2021). Beyond casual resemblance: rigorous frameworks for comparing regeneration across species. Annual Review of Cell and Developmental Biology, 37(1), 415–440.

Srivastava, M., Mazza-Curll, K. L., van Wolfswinkel, J. C., & Reddien, P. W. (2014). Whole-body acoel regeneration is controlled by Wnt and Bmp-Admp signaling. Current Biology, 24(10), 1107–1113.

Stückemann, T., Cleland, J. P., Werner, S., Thi-Kim Vu, H., Bayersdorf, R., Liu, S.-Y., Friedrich, B., Jülicher, F., & Rink, J. C. (2017). Antagonistic Self-Organizing Patterning Systems Control Maintenance and Regeneration of the Anteroposterior Axis in Planarians. Developmental Cell, 40(3), 248–263.e244. 10.1016/j.devcel.2016.12.024

Tratkiewicz, K., & Gąsiorowski, L. (2025). Spontaneous ectopic head formation enables reversal of the body axis polarity in microscopic flatworms. Proceedings of the Royal Society B: Biological Sciences, 292(2057).

True, J. R., & Haag, E. S. (2001). Developmental system drift and flexibility in evolutionary trajectories. Evolution & Development, 3(2), 109–119.

Umesono, Y., Tasaki, J., Nishimura, K., Inoue, T., & Agata, K. (2011). Regeneration in an evolutionarily primitive brain–the planarian *Dugesia japonica* model. European Journal of Neuroscience, 34(6), 863–869.

Umesono, Y., Tasaki, J., Nishimura, Y., Hrouda, M., Kawaguchi, E., Yazawa, S., Nishimura, O., Hosoda, K., Inoue, T., & Agata, K. (2013). The molecular logic for planarian regeneration along the anterior–posterior axis. Nature, 500(7460), 73–76. 10.1038/nature12359

Van Cleave, C. D. (1929). An experimental study of fission and reconstitution in Stenostomum. Physiological Zoology, 2(1), 18–58.

Vellutini, B. C., Martín-Durán, J. M., Børve, A., & Hejnol, A. (2024). Combinatorial Wnt signaling landscape during brachiopod anteroposterior patterning. BMC biology, 22(1), 212.

Vila-Farré, M., Rozanski, A., Ivanković, M., Cleland, J., Brand, J. N., Thalen, F., Grohme, M. A., von Kannen, S., Grosbusch, A. L., & Vu, H. T.-K. (2023). Evolutionary dynamics of whole-body regeneration across planarian flatworms. Nature ecology & evolution, 7(12), 2108–2124.

Wenemoser, D., Lapan, S. W., Wilkinson, A. W., Bell, G. W., & Reddien, P. W. (2012). A molecular wound response program associated with regeneration initiation in planarians. Genes & development, 26(9), 988–1002.

Wiberg, R. A. W., Brand, J. N., Viktorin, G., Mitchell, J. O., Beisel, C., & Schärer, L. (2023). Genome assemblies of the simultaneously hermaphroditic flatworms *Macrostomum cliftonense* and *Macrostomum hystrix*. G3 Genes|Genomes|Genetics, 13(9). 10.1093/g3journal/jkad149

Witchley, Jessica N., Mayer, M., Wagner, Daniel E., Owen, Jared H., & Reddien, Peter W. (2013). Muscle Cells Provide Instructions for Planarian Regeneration. Cell reports, 4(4), 633–641. 10.1016/j.celrep.2013.07.022

Wood, D. E., Lu, J., & Langmead, B. (2019). Improved metagenomic analysis with Kraken 2. Genome Biology, 20(1), 257.

Yazawa, S., Umesono, Y., Hayashi, T., Tarui, H., & Agata, K. (2009). Planarian Hedgehog/Patched establishes anterior–posterior polarity by regulating Wnt signaling. Proceedings of the National Academy of Sciences, 106(52), 22329–22334.

