## Supplementary figures and images for "Divergent evolution of the Wnt signaling system in flatworms"

### Fig. S1

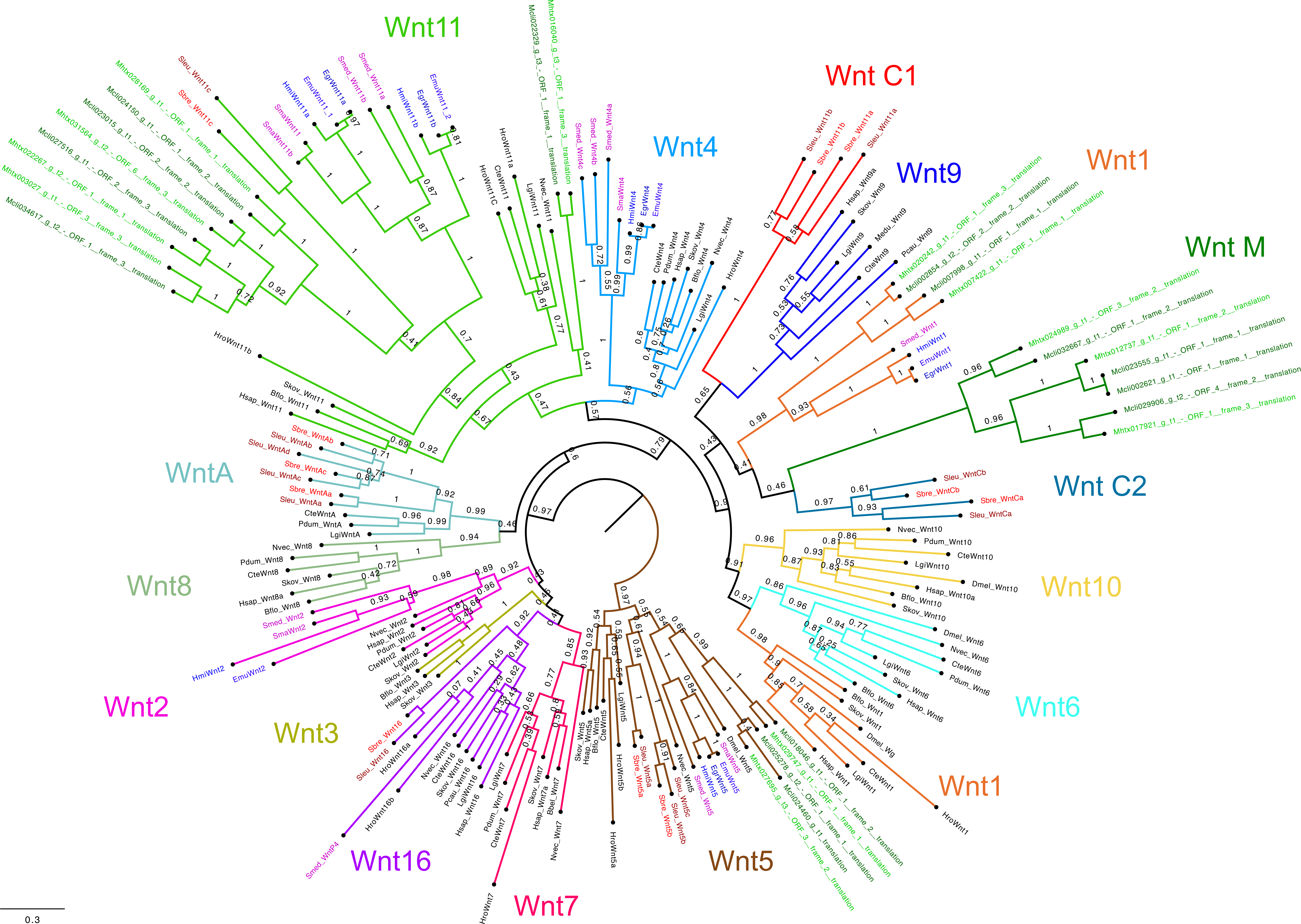

### Fig. S2

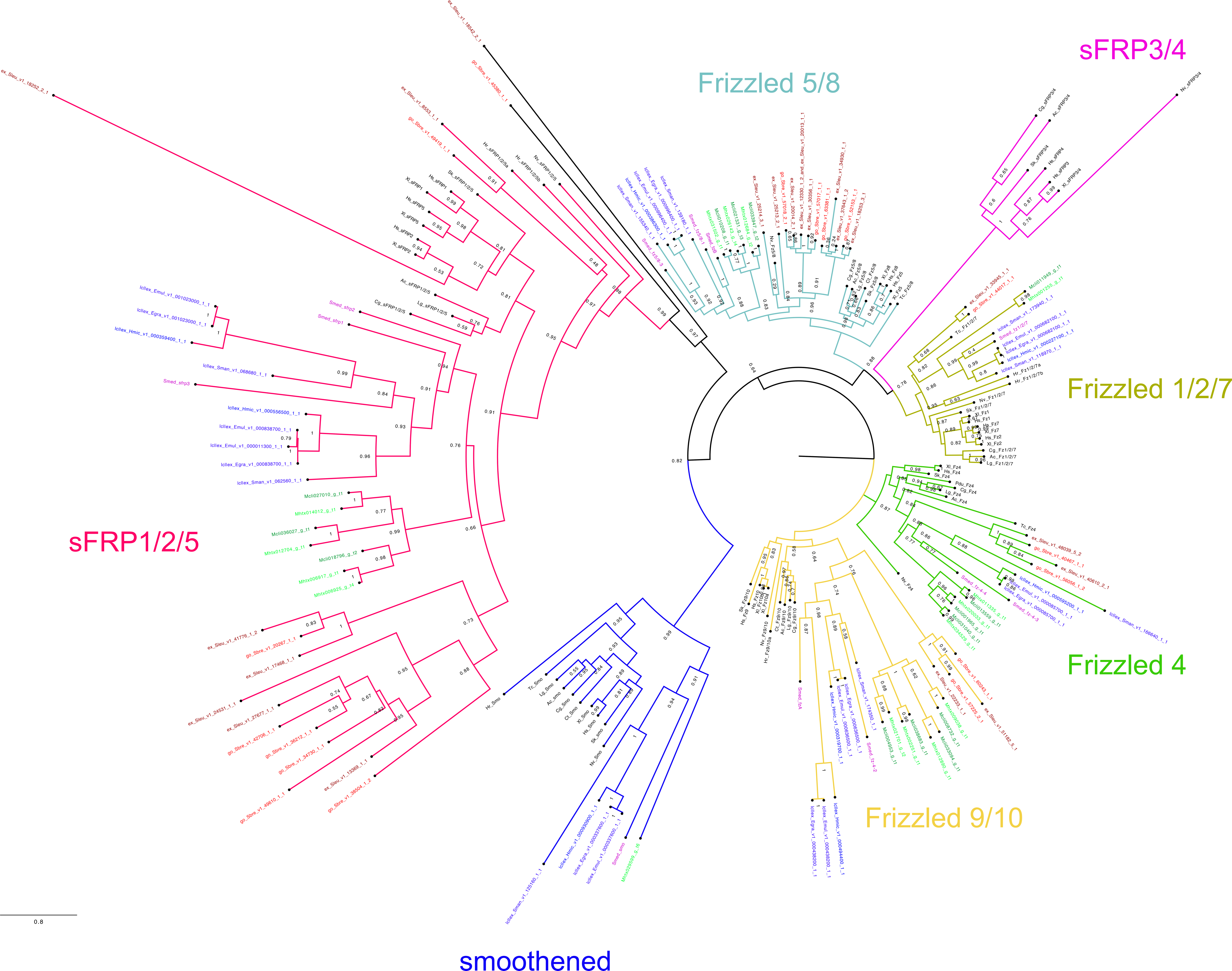

### Fig. S3

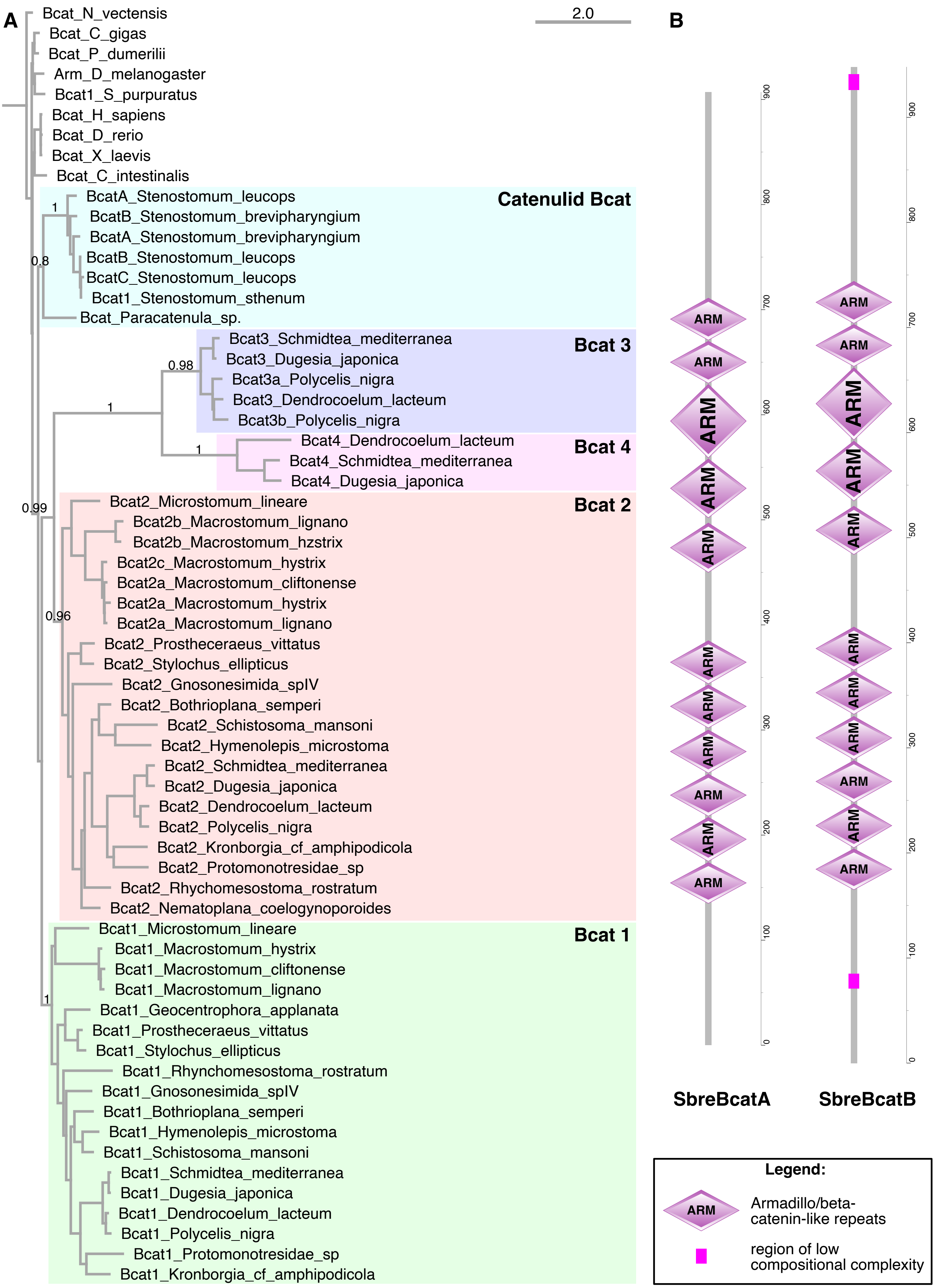
